# Discovery and characterization of bacterial unspecific peroxygenase-like heme-thiolate enzymes

**DOI:** 10.1101/2025.11.07.687166

**Authors:** Esteban Lopez-Tavera, Anton A. Stepnov, Nikolai S. Ersdal, Marta Barros-Reguera, Ronja Marlonsdotter Sandholm, Sabina Leanti La Rosa, Morten Sørlie, Vincent G. H. Eijsink, Gustav Vaaje-Kolstad

**Affiliations:** Faculty of Chemistry, Biotechnology and Food Science, Norwegian University of Life Sciences (NMBU), P.O. Box 5003, N-1432 Aas, Norway

**Keywords:** Peroxygenase, peroxidase, heme-thiolate enzyme, bacterial enzyme, structure database mining, hydrogen peroxide

## Abstract

Unspecific peroxygenases (UPOs, EC. 1.11.2.1) are promising biocatalysts for the oxyfunctionalization of organic molecules and the synthesis of industrially relevant compounds due to their vast repertoire of catalyzed reactions. To date, thousands of putative UPO genes have been identified in eukaryotic genomes, most of them in the *Ascomycota* and *Basidiomycota* phyla, and several UPOs have been characterized. Remarkably, no related enzymes have ever been reported in prokaryotic organisms. Here, we describe the discovery of a novel family of diverse bacterial heme-thiolate peroxygenases through structure database mining, followed by functional characterization of selected representatives. The bacterial UPO-like proteins (BUPOs) are structurally analogous to family I (“short”) fungal UPOs, despite having sequence similarity below 20%. Expression of one of these proteins (*Hyd*BUPO) in its native host (*Hydrogenophaga* sp. A37) was confirmed by proteomics. Several BUPOs were cloned and expressed in *Escherichia coli*. In biochemical assays, the BUPOs were able to catalyze one-electron oxidation (peroxidase activity) of ABTS and 2,6-dimethoxyphenol, and two-electron oxidation (peroxygenase activity) of naphthalene, indole, 3-phenyl-1-propanol and 16-hydroxypalmitic acid, using hydrogen peroxide as co-substrate. These enzymes thus represent a new family of bacterial heme-thiolate peroxygenases that share structural and functional features with eukaryotic UPOs, offering new potential candidates for developing industrially relevant biocatalysts.

## Introduction

Since their discovery in 2004 (1), unspecific peroxygenases (UPOs, EC 1.11.2.1) have been shown to catalyze a remarkably large set of oxyfunctionalization reactions in a broad spectrum of substrates (2, 3). Among the UPO-catalyzed reactions are the oxidation of aryl alcohols to aldehydes and carboxylic acids (1), hydroxylation of aromatic substrates such as toluene and naphthalene (4), oxidative cleavage of ether bonds (5), oxidation of saturated alkanes, aliphatic alcohols and fatty acids to alcohols, ketones and carboxylic acids (6–8), and epoxidation of unsaturated compounds (9). Unlike cytochrome P450 monooxygenases (CYP450), capable of similar oxyfunctionalization reactions, UPOs do not require cofactors such as NAD(P)H or flavin-containing redox partners for catalysis but depend solely on hydrogen peroxide as co-substrate (1).The vast repertoire of peroxygenation reactions catalyzed by UPOs, together with the simple reaction setups requiring only H_2_O_2_, makes this enzyme family attractive for industrial applications. However, despite recent advances in optimizing UPO production (10–12), low expression yields still represent a major bottleneck for the use of UPOs in industrial applications (2, 13).

When UPOs were first described, no sequence homology was found with any previously known peroxidases or cytochrome P450 peroxygenases, and the only known relative was a heme chloroperoxidase from the filamentous fungus *Caldariomyces fumago* (*Cfu*CPO) (14). Since then, mining of genomic databases has revealed thousands of putative UPO genes in several fungal phyla (mainly *Ascomycota* and *Basidiomycota*) and *Oomycota* (13), classified in two large families, I “short” (29 kDa in average), or II “long” (44 kDa in average) (15), several of which have been studied and shown to encode enzymes with a wide diversity of substrates, catalytic activities and stereoselectivities (16, 17). For over twenty years, these enzymes have been considered exclusive to fungi and *Oomycota*, with no confirmed representation in other eukaryotes or any prokaryotes.

Owing to the development of the powerful structure-based search engine FoldSeek (18) and the availability of vast databases of accurate protein structure models such as the AlphaFold database (over 200 million proteins) (19, 20), it is now possible to go beyond sequence similarity when searching for functionally related proteins. Furthermore, in the case of enzymes that depend on cofactors, like heme in UPOs, the availability of tools such as AlphaFold3 (21) for accurate prediction of cofactor binding greatly enhances the capabilities of in silico mining of novel proteins.

Here, we report the discovery of a new family of bacterial UPO-like enzymes, referred to as BUPOs below, characterized by a thiolate-coordinated heme cofactor and a tertiary structure resembling that of eukaryotic UPOs. We verified the expression of one of these enzymes in its native organism and then successfully produced several BUPOs in *Escherichia coli* and demonstrated their UPO-like enzymatic activity. These findings significantly expand the known redox enzyme repertoire of prokaryotes and reveal potential candidates for the development of industrial biocatalysts.

## Results

### 1. Structure-based database mining reveals bacterial UPO-like heme-thiolate proteins

The development of high-precision structure prediction tools such as Alphafold (19) has revolutionized in silico enzyme discovery by allowing structure-based database searches rather than sequence-based searches. To identify putative BUPOs, two crystal structures of family I (“short”) fungal UPOs, *Mro*UPO (PDB 7ZBP) and *Hsp*UPO (PDB 7O1R), and one family II (“long”) member, *Aae*UPO’s evolved mutant PaDa-I (PDB 5OXU), were used as templates for searching similar structures using the FoldSeek online server (18), applying a taxonomic filter for bacteria (eubacteria). While no relevant hits were found for the long UPO, the search with short UPOs yielded a total of 21 bacterial proteins with high probability score (> 0.7) and sequence coverage (> 70 %, relative to the queried proteins). Despite the structural similarity, these proteins shared low sequence identity (< 18 %) with the queried fungal UPOs, which is most likely the reason why bacterial UPO-like proteins have remained undetected by sequence-based search algorithms.

Prediction of the structures of these proteins with a b-type heme cofactor, using the Alphafold 3 server (21) showed a cysteine residue coordinating the heme in the same way as in fungal UPOs, adding to the notion that these proteins indeed are BUPOs. Two out of the 21 proteins (UniProt accession numbers A0A4Q5XIQ3 and A0A2H9SVS9) had a histidine-coordinated heme and were excluded from further analyses.

To expand the dataset, the 19 bacterial sequences were searched in UniProt BLAST towards the UniProtKB database, which resulted in 169 additional proteins with sequence identities varying from 33 to 98 %, relative to the query sequences. The final set of putative bacterial UPOs consisted of 188 unique proteins (**Supplementary Dataset 1**). Then, to decrease redundancy in subsequent analyses, the sequences were clustered using Cd-hit (22), keeping representative sequences with < 90 % sequence identity. This reduced set was used to build a multiple sequence alignment (MSA) with 25 previously studied fungal UPOs (**Table S1**) using the Mafft server (23).

The final alignment, containing 85 bacterial and 25 fungal sequences, was used to build a neighbor-joining phylogenetic tree of UPOs and BUPOs (**Fig. 1**), which shows that the BUPO sequences cluster separately from both long and short fungal UPOs. The tree reflects the considerable diversity among the bacterial sequences. Clade I consists mostly of proteins with a lipoprotein signal peptide (Sec/SPII-type), according to prediction by the SignalP algorithm (24). In clade II, most proteins have a predicted Sec/SPI-type secretion signal peptide, indicating that they are exported from the cytoplasm. While most of the BUPO domains were not assigned a protein family by InterPro Scan (25), the proteins in clade III could be assigned to the chloroperoxidase-like superfamily (ID IPR036851). Not surprisingly, clade III members are also the ones with the highest sequence identity (up to 19%) with fungal UPOs (**Fig. S1**). Several clade III proteins have predicted N-terminal disordered regions. Proteins in clades IV and V have no obvious representative features recognized by InterPro. Finally, whereas clade I-V are single-domain proteins, clade VI has several members that consist of two domains: the N-terminal UPO-like domain (observed in predicted structures; not identified by InterPro), followed by a transmembrane region and a C-terminal di-heme cytochrome C peroxidase domain (InterPro code IPR047758).

**Fig. 1.**
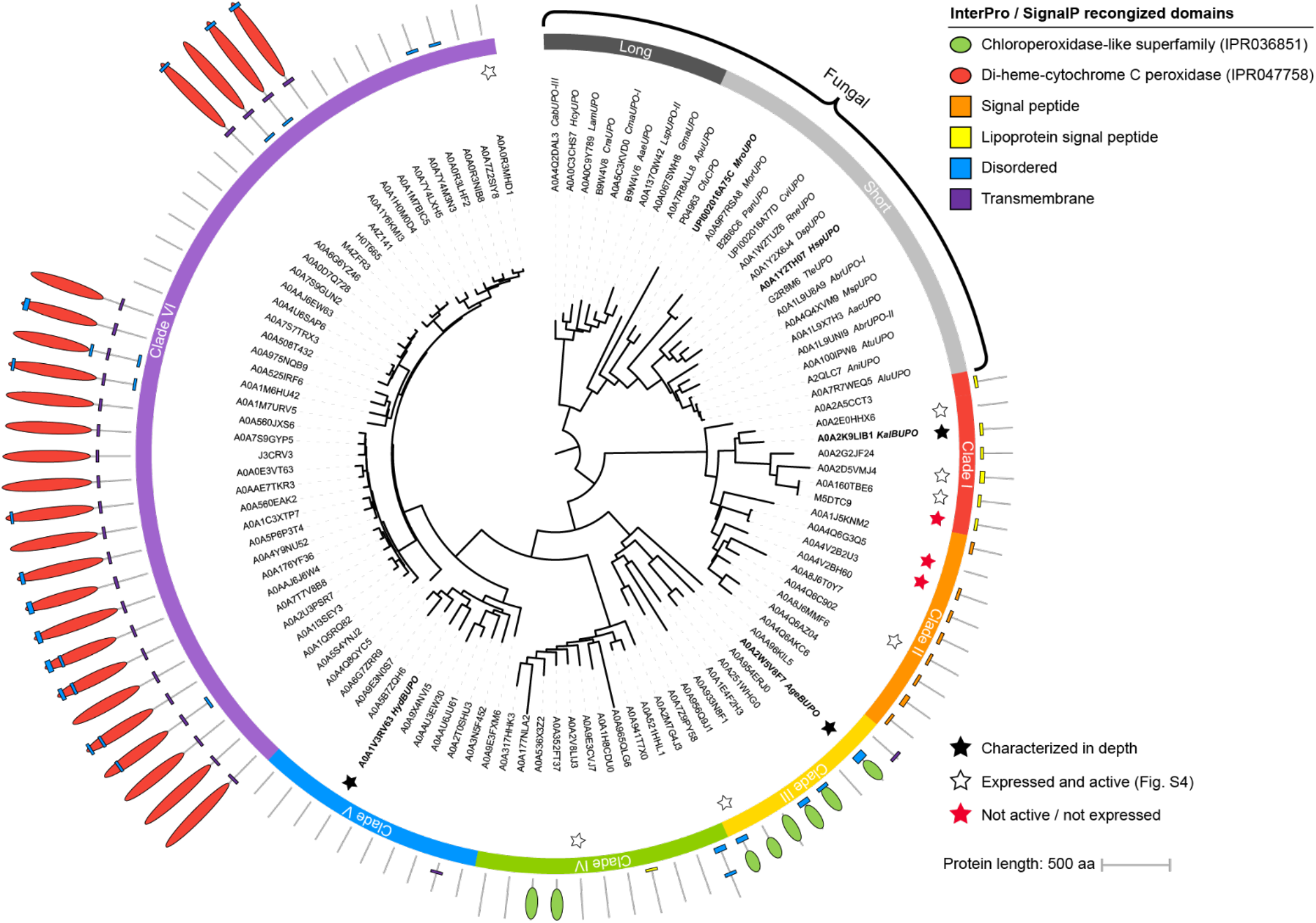
Phylogenetic tree of representative fungal and bacterial UPOs. The phylogenetic tree features 25 previously studied fungal UPOs (family I “short”, light grey; family II “long”, dark grey), and 85 putative bacterial UPOs, separated into 6 distinct clades I-VI. The stars indicate sequences that were tested for expression in *E. coli* in this study, the black stars (★) indicate the proteins that were characterized in depth (see main text), the white stars (☆) indicate the proteins with activity observed in rapid microtiter plate assay, and the red stars 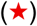 indicate proteins with no detected activity / expression (**Fig. S6**). Protein domains assigned using InterPro and signal peptides by SignalP are shown around the tree, with the N-terminus pointing toward the center of the tree. The protein labeled A0A2A5CCT3 is not part of Clade I but is closely related and is predicted to be a lipoprotein as most of the Clade I members.

Analysis of regions containing key catalytic UPO motifs (**Fig. S2**) revealed that the PCP motif, which includes the heme-coordinating cysteine (Cys39 in *Hsp*UPO), is conserved among the BUPOs, as well as an acid-base catalytic pair (Glu180 as acid-base catalyst and His110 as its charge stabilizer in *Hsp*UPO) that is known to participate in the heterolytic cleavage of H_2_O_2_ (13). In some of the bacterial proteins, His110 is replaced by another basic residue (lysine or arginine). Of note, arginine is also observed in the acid-base pair in long UPOs (13). In contrast, residues Glu109, Asp111 and Ser113 in *Hsp*UPO, involved in binding of a Mg^2+^ ion and highly conserved in fungal UPOs (13), seem to be completely absent in the bacterial proteins (**Fig. S2 B**). In fungal UPOs, the bound magnesium ion is thought to stabilize the strained porphyrin system of the heme cofactor (13).

### 2. The bacterial UPO-like proteins feature key structural elements of fungal UPOs

To couple the sequence analysis to structural information, we predicted the structures of the 85 putative BUPOs shown in **Fig. 1**, using the Alphafold 3 server (21), including a b-type heme cofactor and a Mg^2+^ ion as ligands. Like the fungal UPOs, BUPOs display a chloroperoxidase-like fold with an orthogonal bundle architecture of mainly alfa-helices. This architecture is exemplified by three BUPOs shown in **Fig. 2 B-D**, which correspond to the enzymes investigated in detail in this study (*Age*BUPO, *Kal*BUPO and *Hyd*BUPO). The TM-scores obtained when comparing *Hsp*UPO (PDB 7o1r) with each of the three predicted BUPO structures shown in **Fig. 2 B-D** varied from 0.66 to 0.74, which is well above the cutoff (0.5) for defining structures as having mostly the same fold (26– 28). The active site of UPOs is characterized by a cysteine coordinating the heme iron at the proximal side and an acid-base pair on the distal side of the heme that is necessary for the cleavage of hydrogen peroxide in the catalytic cycle (29). All these elements are present in BUPOs (**Fig. 2 F-H**), and, as mentioned above, these residues are highly conserved among all putative bacterial UPOs (**Fig. S2**). Note that the histidine in the conserved acid-base pair (His110 & Glu180 in *Hsp*UPO) is replaced by another basic residue (Lys95) in *Hyd*BUPO (**Fig. 2 H**). Notably, magnesium binding was predicted in a different loop for *Age*BUPO and *Kal*BUPO (**Fig. 2 B** and **C**) than in fungal UPOs. While the magnesium ion is predicted to bind in *Hyd*BUPO (**Fig. 2 D** and **H**) in an analogous location to that of fungal UPOs, later analysis by ICP-MS showed that this protein does not bind Mg^2+^ (see below).

**Fig. 2.**
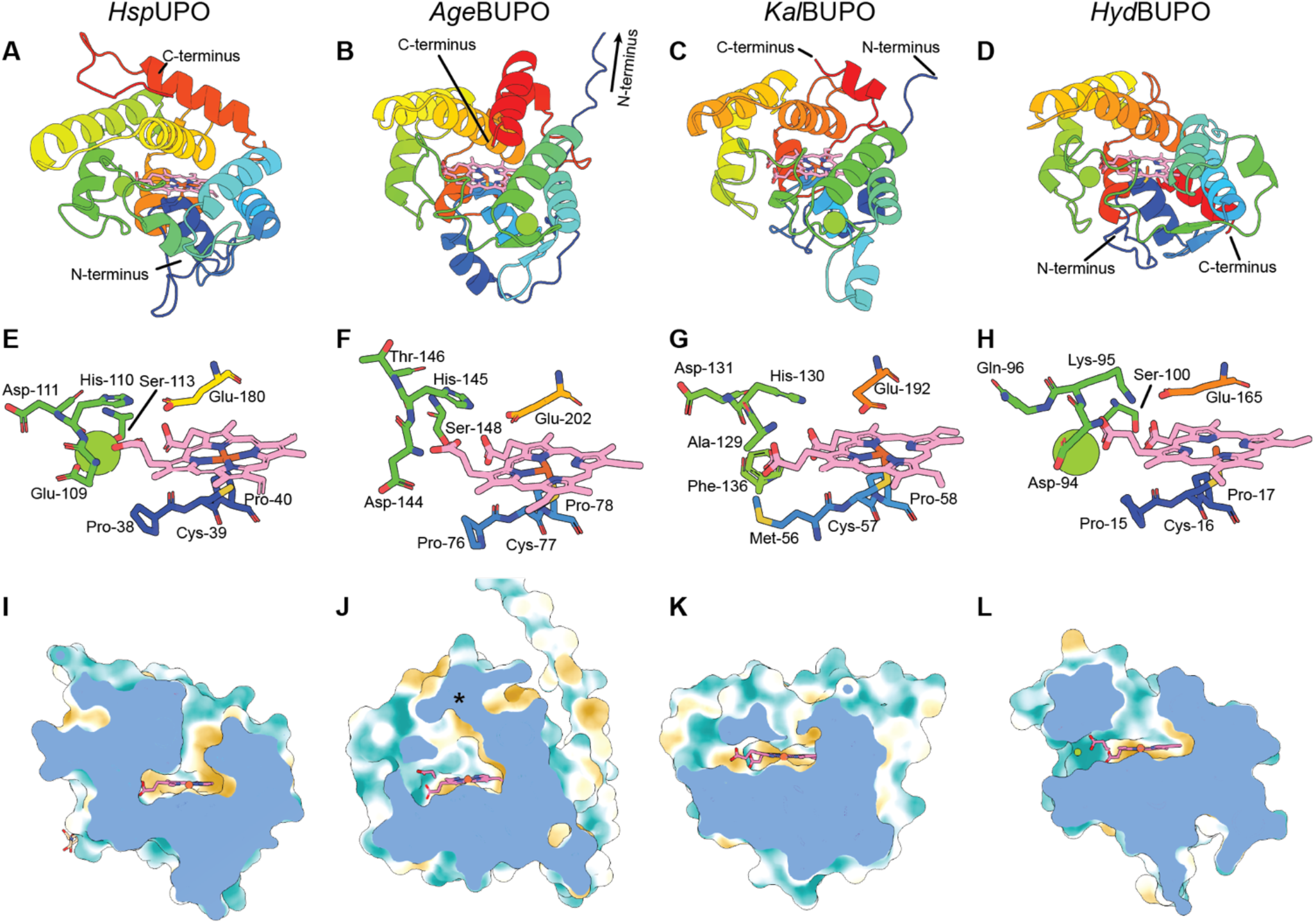
Structural features of bacterial UPOs and comparison with *Hsp*UPO. The short fungal UPO *Hsp*UPO (PDB 7o1r; **A, E, I**) is shown as reference for comparison with the AlphaFold3-predicted structures of *Age*BUPO (**B, F, J**, clade 3), *Kal*BUPO (**C, G, K**, clade 1) and *Hyd*BUPO (**D, H, L**, clade 5). In the top row (**A-D**), the structures are colored by residue position to illustrate the topology and spatial arrangement of the α-helices. The UPO-defining conserved residues are shown in the second row (**E-H**), together with the heme group and van der Waals sphere of Mg^2+^ (green). The residue numbering is relative to the full peptide chain, including signal peptides. The solvent excluded surface (**I-L**) is shown in a lipophilicity color scale, from gold (lipophilic) to teal (hydrophilic). The transversal view of the access tunnel (**I-L**) shows a second opening present in the BUPOs (extending to the left) where one of the heme propionate groups is exposed to the solvent. The C-terminus of *Age*BUPO shown in panels **B** and **J** (marked in **J** with a star symbol, **\***), as well as a long N-terminal region (truncated in the figure), have a very low prediction confidence (pLDDT value < 50) and are likely to be misfolded in the model or inherently disordered. The Alphafold3 models colored by pLDDT score are shown in **Fig. S4**.

The predicted structures suggest diverse geometries and sizes for the active site access tunnels of the bacterial UPOs (**Fig. 2 J-L**). Whereas fungal UPOs only have one tunnel leading to the active site (e. g. *Hsp*UPO, **Fig. 2 I**), all BUPOs except those in clade IV are predicted to have a second tunnel leading to the heme cofactor, where one of the heme propionate groups is exposed to the solvent (**Fig. 2 F-H**, and **Fig. S3**). Notably, the location of the second tunnel entrance in BUPOs is similar to the location of the Mg^2+^ coordination site in fungal UPOs (**Fig. 2, E-L**).

### 3. The native expression of a BUPO is confirmed in *Hydrogenophaga sp*. A37 through proteomics

Having identified BUPO candidates in sequence databases, we sought to experimentally confirm the presence of one such gene in its native organism’s genome and to determine whether the corresponding BUPO was natively expressed, in order to rule out the remote possibility that these bacterial genes represent sequencing artifacts or non-functional evolutionary remnants. For this purpose, we used *Hydrogenophaga sp*. A37, a denitrifying bacterium isolated from soil (30), which harbors the BUPO candidate with UniProt accession A0A1V3RV63 (clade V), hereafter referred to as ***Hyd*BUPO**. At the start of this study, the genome assembly of this organism available in NCBI (GCA_002001205.1) had been generated using short-read Illumina sequencing. It comprised 195 contigs with an N50 of 56 kb and a total genome size of 5.6 Mb. Here, we generated an improved *Hydrogenophaga* sp. A37 genome assembly using Oxford Nanopore long-read sequencing (GCA_053010425.1). The new assembly consists of a single circular chromosome of 5,605,460 bp (assembly level: complete genome), with an overall G+C content of 65.5%. The genome achieved 80 × coverage. Genome quality assessments indicated 99.1% completeness and 1.9% contamination. In total, 5295 CDS, 5226 genes, 45 tRNAs, and 3 complete rRNA were predicted. Sequence alignment using Minimap2 (31) and DGENIES (32) revealed 99.86 % identity with the earlier genome version, indicating high sequence identity and collinearity (**Fig. S5**). With this improved genome, we generated a complete proteome to be used as the reference database for the proteomic analysis of *Hydrogenophaga sp*. A37. The bacterium was grown for 48 h at 22 °C in a diluted commercial medium (tryptone soy broth), and proteins were isolated from the cell pellet prior to trypsin digestion and LC-MS/MS analysis. A total of 2,179 proteins were detected (listed in **Supplementary Dataset 2**), among which we identified *Hyd*BUPO (NCBI accession MGS5087929.1), thereby confirming that *Hyd*BUPO is a functional gene expressed in its native organism.

### 4. Functional bacterial UPO-like enzymes can be expressed in *Escherichia coli*

A set of 13 candidate BUPOs spanning the six different clades (**Fig. 1**) were selected for heterologous expression in *Escherichia coli*. For the 9 candidates with predicted signal peptides or N-terminal disordered regions, genes encoding three variants were synthesized: one with the native signal peptide, one with the pelB signal peptide, and a truncated variant with no signal peptide (see **Supp M&M** and **Table S2** for details). For the remaining four candidates, only the native variant was synthesized. A micro-scale expression screen based on detecting activity towards 2,2’-azino-bis(3-ethylbenzothiazoline-6-sulfonic acid (ABTS) indicated that 10 of the 13 functional BUPOs were successfully produced, with at least one enzyme variant showing peroxidase activity towards ABTS (**Fig. S6 B**). Guided by expression levels and diversity criteria (i.e., including representatives from each of the most distinct clades), three enzymes were selected for further characterization: *Age*BUPO (BU04-1) from *Archangium gephyra* (clade III, UniProt accession A0A2W5V8F7), *Kal*BUPO (BU06-2) from *Ketobacter alkanivorans* (clade I, A0A2K9LIB1), and *Hyd*BUPO (BU12-1) from *Hydrogenophaga sp*. A37 (clade V, A0A1V3RV63). All three proteins were purified to homogeneity (**Fig. S7**) yielding 40 mg, 10 mg and 10 mg of heme-loaded enzyme per liter of culture (*Age*BUPO, *Kal*BUPO and *Hyd*BUPO, respectively), based on absorbance at 420 nm (Soret band). This represents, to the best of our knowledge, the highest yield reported to date for heme-loaded soluble UPOs expressed in *E. coli* (33) which is, perhaps, expected, given the bacterial origin of these enzymes.

After purification, the UV-Vis spectra of the resting state and of the dithionite-reduced proteins were measured. In the resting state, the BUPOs feature the characteristic Soret band with maxima at 423 nm for *Age*BUPO and *Kal*BUPO, and 425 nm for *Hyd*BUPO, as well as additional peaks at 355-359 nm, 542-544 nm, and 576-580 nm (**Fig. S8**). Upon reduction with sodium dithionite, the Soret maxima shifted to 400 nm (*Age*BUPO), 412 nm (*Kal*BUPO) and 402 nm (*Hyd*BUPO), similar to what has been reported for fungal UPOs (1, 34).

To confirm the amounts of heme-loaded enzyme and investigate the possible presence of a magnesium binding site in the BUPOs, the iron and magnesium contents were measured through inductively coupled plasma mass spectrometry (ICP-MS). Based on the determined iron contents, the BUPO extinction coefficients at the Soret maxima were determined to be 101.62, 106.57, and 110.05 mM^-1^ cm^-1^ for *Age*BUPO, *Kal*BUPO and *Hyd*BUPO, respectively. Interestingly, the magnesium content was negligible in all three BUPOs, whereas iron and magnesium were almost equimolar in *Hsp*UPO (**Fig. S9**). In fungal UPOs, magnesium contributes to stabilizing the strained heme cofactor (13). The negligible magnesium content in BUPOs thus points to an alternative evolutionary solution for maintaining heme stability in bacterial enzymes.

### 5. BUPOs are capable of one- and two-electron oxidations of aromatic substrates

Given the active site similarities observed between fungal UPOs and BUPOs, it was of interest to determine whether these bacterial proteins were capable of both one-(peroxidase) and two-electron (peroxygenase) oxidations of typical chromogenic UPO substrates. Indeed, the rapid functional screen had already indicated peroxidase activity towards ABTS (**Fig. S6**). Peroxidase activity of the three selected purified BUPOs was further explored using more controlled conditions with both ABTS and 2,6-dimethoxyphenol (2,6-DMP) as substrates, where the latter dimerizes to produce coerulignone after one-electron oxidation (35). All BUPOs showed peroxidase activity with at least one of the substrates, with considerable variation among the enzymes (**Fig. 3, Fig. S10**). 0.5 µM *Age*BUPO and *Kal*BUPO showed initial rates of 5.2 and 5.9 µM min^-1^, respectively, in reactions with 0.8 mM ABTS at pH 5.5 (**Fig. 3**). *Hyd*BUPO, on the other hand showed little activity towards this substrate under these conditions, exhibiting only marginally higher rate compared to hemin chloride at the same concentration (0.5 µM). Under these conditions, the fungal enzyme *Hsp*UPO, used as a positive control, was approximately four times faster when used at 10-fold lower concentration (0.05 µM; initial rate of 21 µM min^-1^). pH-activity profiles showed that the pH optimum for oxidation of ABTS by *Hyd*BUPO is at pH 7.0, whereas for *Age*BUPO and *Kal*BUPO the optimum pH is at 5.6 and 5.0, respectively (**Fig. S11**). Even at its optimum pH, *Hyd*BUPO reached less than half the rate of *Age*BUPO and *Kal*BUPO. The pH-activity profiles confirmed the superior performance of *Hsp*UPO on ABTS. In the 2,6-DMP assay, carried out at pH 6.0 with 0.5 µM BUPO, *Kal*UPO showed the highest rate (7.6 µM min^-1^), followed by *Hyd*BUPO (4.5 µM min^-1^), while *Age*BUPO reached only 1.3 µM min^-1^ (**Fig. 3**). Again, *Hsp*UPO outperformed the BUPOs (75 µM min^-1^, using 0.05 µM enzyme).

**Fig. 3.**
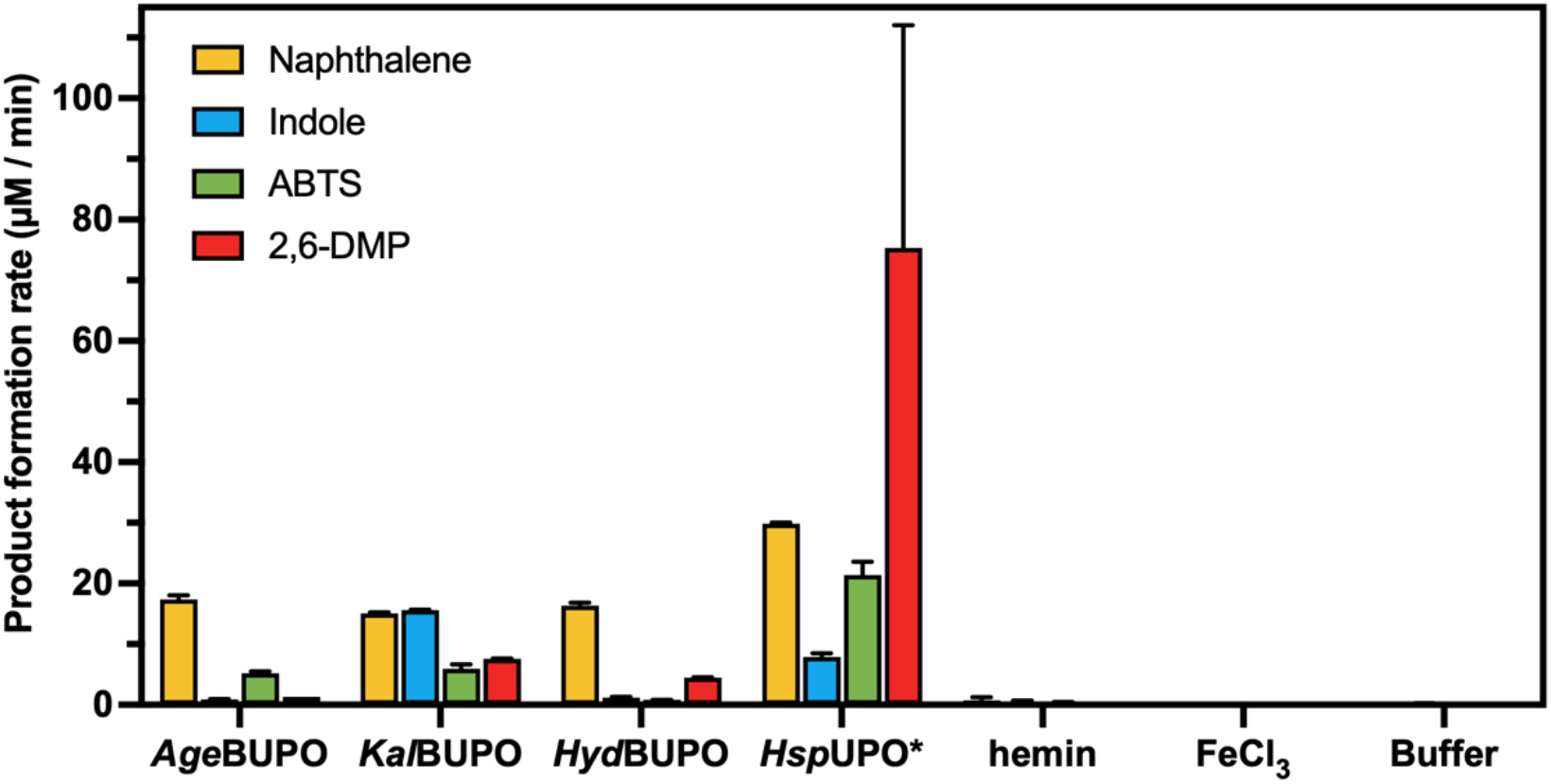
Initial rates for the oxidation of four model substrates by *Age*BUPO, *Kal*BUPO, *Hyd*BUPO and *Hsp*UPO. The peroxidase reactions on ABTS and 2,6-DMP were measured in 96-well plates using 0.5 µM enzyme (0.05 µM for *Hsp*UPO), and the peroxygenase reactions on naphthalene and indole were measured using 5 µM enzyme (0.5 µM for *Hsp*UPO). The formation of oxidized products was measured for 15 min at the following wavelengths: 2,6-DMP, 469 nm (ε_469_ = 27.5 mM^−1^ cm^−1^ (35)); ABTS, 418 nm (ε_418_ = 36 mM^−1^ cm^−1^ (35)); indole, 670 nm (ε_670_ = 4.8 mM^−1^ cm^−1^ (35)); naphthalene, 324 nm (quantified with a standard curve of 1-naphthol). For further details on the reaction conditions, see **Supplementary M&M**. The reaction traces are shown in **Fig. S10**. The error bars indicate the standard deviation of replicates (n=3). **\***Note that *Hsp*UPO was used at 10-fold lower concentration than the rest of the enzymes in all reactions.

Catalyzing peroxygenase reactions is what makes UPOs interesting for producing industrially relevant molecules (13). Two examples of such molecules are 1-naphthol, which is a precursor for herbicides, insecticides and pharmaceuticals (36), and indigo, which is a widely used dye (35). These two compounds are also used as colorimetric indicators of peroxygenase reactions, since naphthalene and indole can be hydroxylated by UPOs to produce 1-naphthol and indigo (upon dimerization of 3-hydroxyindole), respectively (35). All three bacterial enzymes showed peroxygenase activity towards naphthalene at pH 5.5, with rates (on the order of 15 µM min^-1^, with 5 µM enzyme) were about ten-fold lower compared to fungal *Hsp*UPO (**Fig. 3**). Regarding the oxidation of indole, only *Kal*BUPO showed clear activity, with a rate (15.6 µM min^-1^ with 5 µM enzyme) that was only ∼5-fold lower than the rate observed for fungal *Hsp*UPO.

Together, these results confirm the ability of BUPOs to catalyze both one- and two-electron oxidation reactions and thus their functional similarity to fungal UPOs.

### 6. BUPOs can oxidize aliphatic substrates

The hallmark feature of unspecific peroxygenases is their ability to perform two-electron oxidations of a broad range of substrates. After observing activity on the aromatic substrates described above, we carried out additional experiments with 3-phenyl-1-propanol (3PP), palmitic acid and 16-hydroxypalmitic acid to assess the capacity of BUPOs to oxidize aliphatic compounds. While 3PP is an aromatic compound, it has an aliphatic moiety which is expected to be oxidized by at least some canonical fungal UPOs (37, 38). Indeed, a control experiment with *Hsp*UPO (**Fig. S12**) showed enzyme-dependent formation of 3-phenylpropionic acid (3PP-A), along with other oxidation products. In comparison, *Kal*BUPO also generated considerable amounts of 3PP-A, while only trace amounts were detected for *Hyd*BUPO and none for *Age*BUPO (**Fig. S12**). Substrate depletion experiments showed similar turnover of 3PP for *Hsp*UPO and *Kal*BUPO after 1 h of reaction, although the fungal enzyme is much faster than the bacterial enzyme (**Fig. 4 A and B**).

**Fig. 4.**
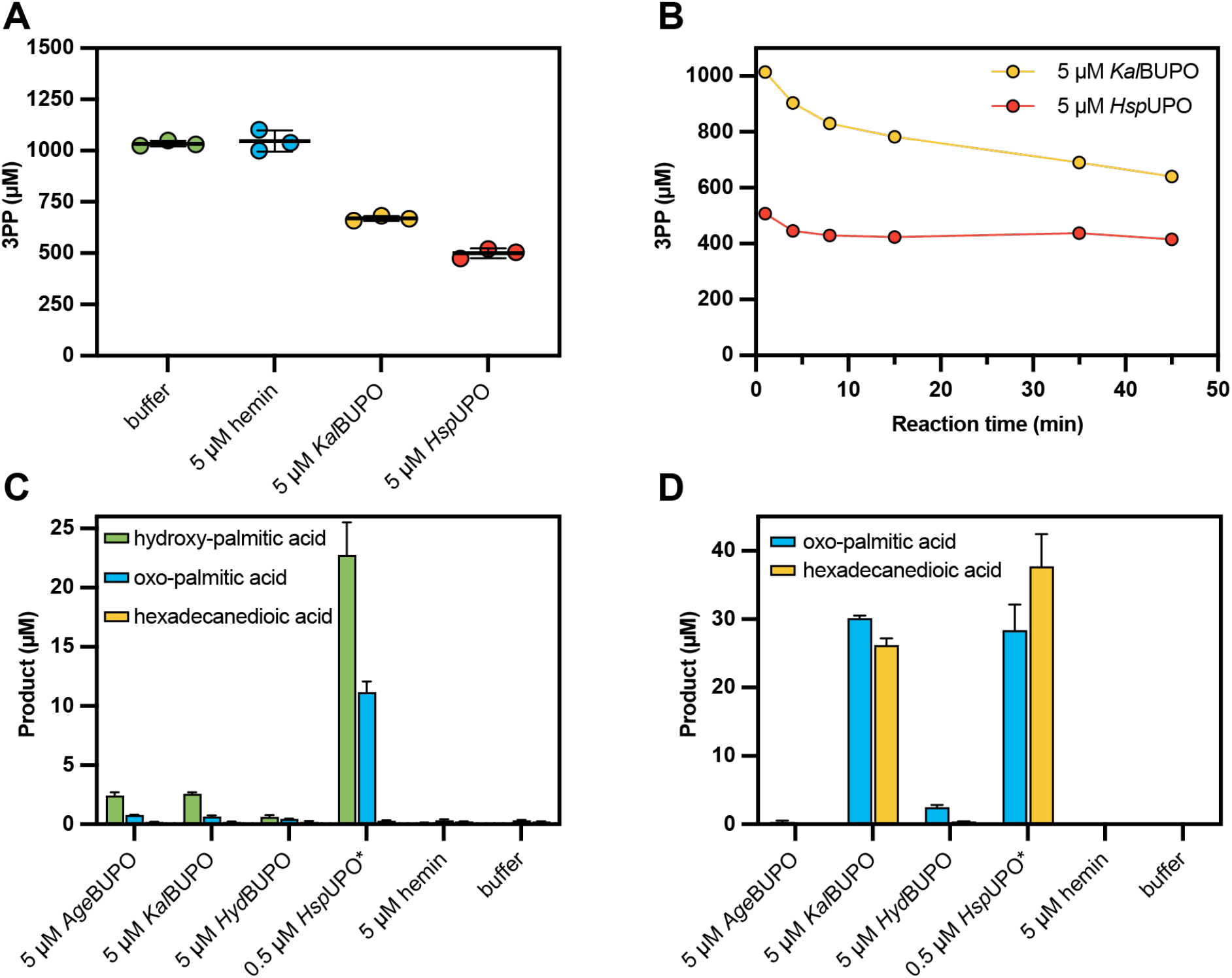
Reactions of BUPOs and *Hsp*UPO with 3-phenyl-1-propanol, palmitic acid and 16-hydroxypalmitic acid. Panel **A** shows substrate depletion in one hour reactions with 3PP, while panel **B** shows reaction time courses for the only two enzymes for which significant 3PP-oxidizing activity was detected, *Kal*BUPO and *Hsp*UPO. The experiments shown in **A** and **B** were carried out in 50 mM sodium phosphate buffer, pH 6.0, at 30 °C (**A**) or at room temperature (**B**). For the time course experiments (**B**), reactions were stopped by addition of catalase (250 u mL^-1^). Panels **C** and **D** show the product formation in reactions with palmitic acid and 16-hydroxypalmitic acid, respectively, carried out with 5 µM BUPO or hemin chloride, or with 0.5 µM *Hsp*UPO, using 100 µM substrate and 1 mM H_2_O_2_ in 10 mM ammonium acetate, pH 6.0, and 20 % acetone. The reactions were incubated for 1 hour at 30 °C and the products were quantified through LC-MS, using the extracted ion chromatograms of the [M-H]^-^ m/z values expected from the hydroxylated products (271.23), aldehyde/ketones (269.21, labeled oxo-palmitic acid), and hydroxy-ketone or dicarboxylic acid (285.21; labeled hexadecanedioic acid). Error bars in panel **A, C** and **D** indicate standard deviations between triplicates, while horizontal lines in **A** denote mean values. Each point in panel **B** is a single measurement. Note that most of the substrate depletion by *Hsp*UPO happened before the first measurement and is not captured in the progress curve.

Encouraged by the activity of *Kal*BUPO towards 3PP, we further explored the activity towards purely aliphatic compounds, using palmitic acid as substrate. In reactions with 100 µM palmitic acid and 5 µM enzyme, only 0.6 - 2.6 µM of hydroxylated product were detected; (**Fig. 4 C**), which translates to less than 1 turnover per enzyme. Nonetheless, these product levels were significantly higher than the level observed in the reactions with hemin chloride instead of enzyme (0.05 ± 0.1 µM). Using the same conditions, 0.5 µM *Hsp*UPO produced 22.8 µM hydroxy and 11.2 µM ketone/aldehyde products (**Fig. 4 C**), which corresponds to approximately 90 turnovers, considering that ketone/aldehyde products are the result of two consecutive oxidations of palmitic acid.

Given that varying amounts of carbonyl products (ketone / aldehyde) were detected in reactions with palmitic acid, we decided to use the terminally hydroxylated compound 16-hydroxypalmitic acid as a substrate (100 µM; **Fig. 4 D**). For this substrate, *Kal*BUPO showed considerably higher activity, with 5 µM enzyme producing 30.2 µM of aldehyde (one two-electron oxidation) and 26.8 µM of hexadecanedioic acid (two two-electron oxidations), which corresponds to 57 % conversion of the substrate, and 17 turnovers per enzyme. Similarly, *Hsp*UPO (0.5 µM) was more active on the hydroxylated substrate, producing 28.4 µM of aldehyde and 37.7 µM of hexadecanedioic acid, which corresponds to 66 % substrate conversion, and 208 turnovers per enzyme. On the other hand, *Age*BUPO showed negligible activity on this substrate, while *Hyd*BUPO produced a total of 2.9 µM products, meaning less than one turnover per enzyme.

Taken together, the results from the 3PP and 16-hydroxypalmitic acid assays indicate that *Kal*BUPO and, to a lesser extent, *Hyd*BUPO can oxidize primary alcohol groups in both aromatic and aliphatic substrates, while marginal activity of the three BUPOs was detected toward unactivated aliphatic C-H bonds in palmitic acid.

### 7. *Kal*BUPO is prone to inactivation by H_2_O_2_

It is not uncommon for redox-active enzymes, including UPOs, to be inactivated due to oxidative damage caused by either co-substrates like H_2_O_2_ or reactive reaction intermediates (39). Notably, the conditions typically used in the laboratory, and in this study, when working with UPOs are quite different from the *in vivo* situation, in which H_2_O_2_ concentrations as high as 1 mM are not likely to occur. To investigate this further, 20 µM *Kal*BUPO was pre-incubated with 1 mM H_2_O_2_ for time intervals varying from 0 to 30 min, before activity was measured using the naphthalene assay. The results showed a decrease in activity that was proportional with the pre-incubation time (**Fig. S13 A**), clearly indicating that the enzyme was inactivated by exposure to H_2_O_2_ in the absence of an oxidizable substrate. Expectedly, measurement of the absorbance at 420 nm of the pre-incubated enzyme showed that the loss of activity was accompanied by heme bleaching (**Fig. S13 B**). It is worth noting that *Kal*BUPO showed a linear progress curve for the reaction with naphthalene, despite the presence of 1 mM H_2_O_2_ (**Fig. S10 C**), which shows that naphthalene is a *bona fide* substrate for this BUPO.

## Discussion

Since their discovery, UPOs have been considered restricted to eukaryotes, largely dominated by fungi, because even sensitive sequence-profile searches failed to detect counterparts in prokaryotes (13). Exploiting the greater evolutionary conservation of 3D structure over primary sequence, we performed structure-based searches seeded with a fungal UPO and identified UPO-like proteins in bacteria. Despite their low sequence identity, the structural similarity between bacterial and fungal UPOs together with the conservation of key active site features suggests divergence from an ancient common ancestor rather than convergence on an identical peroxygenase architecture.

The hallmark of UPO catalysis is H_2_O_2_-driven oxyfunctionalization, i.e. insertion of oxygen into C– H / C=C bonds, alongside one-electron (peroxidase-type) oxidations across diverse substrates. The three BUPOs characterized in this work fulfill these criteria: they oxidize canonical UPO substrates (e.g., ABTS and naphthalene) as well as other compounds (e.g., 3-phenyl-1-propanol, palmitic acid and 16-hydroxypalmitic acid), using H_2_O_2_ as the co-substrate. Compared to the reference fungal UPO, apparent reaction rates are generally lower. Whether this reflects true catalytic differences or suboptimal assay conditions remains unclear, as the biologically relevant substrates of BUPOs, and of many fungal UPOs, are still unknown.

An intriguing difference that separates the BUPOs from fungal UPOs is the presence of a second access channel to the heme, which exposes one of the heme propionate groups to the solvent. In fungal UPOs, this same propionate group participates in coordinating the Mg^2+^ ion that helps stabilize the strained porphyrin system (13), suggesting that the bacterial enzymes may have evolved a distinct structural solution, with potential functional implications. Comparable multi-tunnel architectures are observed in other heme enzymes, including dye-decolorizing peroxidases (DyPs) and cytochrome P450s. In DyPs, two main channels lead to opposite sides of the heme, and a third exposes a propionate to solvent (40), closely resembling the secondary channel identified in BUPOs. Although the role of this channel in DyPs remains uncertain, it has been proposed to accommodate bulky aromatic substrates such as monolignols for oxidation. In cytochrome P450s, multiple dynamic channels have been described, providing pathways for substrates, co-substrates, water, and protons (41). By analogy, the presence of two channels in BUPOs may enable complementary functions, one serving as an entry route for the co-substrate and the other facilitating electron transfer from the main substrate to the heme. This architectural divergence from fungal UPOs may therefore reflect a bacterial adaptation toward broader substrate access or different dynamics of (co-)substrate flowing in and out of the active site.

Given the pronounced structural differences between BUPOs and fungal UPOs, it is not surprising that their reactivity profiles also diverge. The BUPOs characterized here catalyze the same types of peroxygenase reactions as fungal UPOs, but generally with lower efficiency. Nonetheless, *Hsp*UPO, used as reference, is among the most efficient members of its family (42) on many typical UPO substrates, which may accentuate the contrast. Exceptions such as *Kal*BUPO, which shows marked activity toward 16-hydroxypalmitic acid, suggest that the natural substrates of these bacterial enzymes differ substantially from the model compounds used here.

While fungal UPOs are highly efficient biocatalysts, their heterologous production remains challenging, often requiring extensive screening and optimization (16). In contrast, bacterial enzymes are typically easier to produce and engineer, and our expression screen showed that 10 of 13 BUPO genes yielded active enzyme without any optimization. The three BUPOs characterized here were produced in a standard *E. coli* strain using only codon optimization and His-tag addition, yielding the highest levels of soluble, heme-loaded enzyme reported for any UPO expressed in bacteria. Thus, BUPOs combine the catalytic versatility of UPOs with the genetic accessibility of bacterial systems, which may facilitate mechanistic studies and open new avenues for peroxygenase engineering.

The physiological roles of BUPOs remain uncertain. Their phylogenetic and structural diversity, and broad but weak activity suggest a range of potential functions. In fungi, UPOs have been suggested to participate in processes such as xenobiotic degradation (15), which is also a plausible role of BUPOs. The relatively low apparent rate of BUPOs with H_2_O_2_ as a co-substrate may indicate that these enzymes prefer alternative co-substrates, such as organic peroxides. Thus, the reactions characterized here could represent side activities, while their main biological function remains to be discovered. Notably, even for fungal UPOs, two decades of research have not fully clarified their native roles.

Sequence analysis revealed a relatively small BUPO family, about 190 sequences compared to over 4,000 fungal UPOs. Despite its limited number, BUPOs are divided in highly distinct clades. Clade I BUPOs, which include *Kal*BUPO, originate from marine bacteria such as *Ketobacter alkanivorans*, an alkane-degrading species harboring *alkB* hydroxylase genes (43). The ability of *Kal*BUPO to oxidize aliphatic substrates like palmitic and 16-hydroxypalmitic acids supports a potential role in alkane metabolism, possibly following initial hydroxylation by AlkB.

On a side note, nearly 70% of all identified BUPO sequences cluster in clade VI, found almost exclusively in *Bradyrhizobium* species. Many of these encode a bi-modular protein containing both a BUPO and a di-heme cytochrome c peroxidase (CCP) domain separated by a transmembrane helix. Bacterial CCPs are known to be ubiquitous proteins whose proposed function is H_2_O_2_ detoxification (44) and in some cases allowing respiration with H_2_O_2_ as the terminal electron acceptor (45); however, a bi-modular architecture where the two modules tethered by a transmembrane linker has never been reported before.

In summary, BUPOs represent a new family of bacterial heme-thiolate enzymes that share the structural fold and motifs of fungal UPOs. Beyond expanding the known catalytic repertoire of bacteria, these findings highlight the power of large-scale structural prediction and genome mining to uncover overlooked enzyme families. Future work should clarify the biological roles of BUPOs, explore their catalytic scope, and harness their bacterial expression advantages for enzyme engineering.

## Materials and Methods

Comprehensive descriptions of experimental procedures, sequence analyses, and analytical methods are provided in the Supplementary Methods.

### Structure-based search and dataset curation

FoldSeek searches (18) were run against AFDB50 using three fungal UPO templates (*Mro*UPO PDB 7zbp, *Hsp*UPO PDB 7o1r, PaDa-I PDB 5OXU) in 3Di/AA mode. Hits were retained with FoldSeek probability > 0.5 and length > 150 aa. The 19 initial candidates were expanded by individual UniProt BLAST searches (E < 10^−18^). Duplicates were removed with CD-HIT (22) at 100% identity, producing the full set of unique sequences, which was reduced with CD-HIT at 90% identity to a representative subset for phylogenetic analysis. Signal peptides and domain annotations were predicted with SignalP 6.0 (24) and InterProScan (25).

### Phylogenetic analysis

Multi-domain proteins were truncated to the UPO-like domain based on AlphaFold models; truncated sequences were aligned with Clustal Omega and then with MAFFT (23) including 25 fungal UPOs. A neighbor-joining tree was built on the MAFFT server using the JTT model with 100 bootstrap replicates; the tree was annotated with iTol (46) together with SignalP/InterPro outputs. Sequence logos were made with WebLogo3 (47). Thirteen putative BUPOs covering the tree diversity were selected for experimental characterization.

### Structure prediction and comparison

AlphaFold3 predictions (21) were generated for the BUPOs including a heme-b cofactor and Mg^2+^. Predicted models were aligned to *Hsp*UPO (PDB 7o1r) using TM-align (27) to obtain TM-scores.

### Materials, gene constructs and expression vectors

Chemicals and reagent sources are listed in the SI. Genes were codon-optimized for *E. coli* and synthesized with N- or C-terminal His tags (**Table S2**). For sequences with predicted signal peptides three variants were made (native, truncated, pelB-SP); lipidation site cysteines were removed where appropriate. Constructs were cloned into pET-29b(+) (NdeI/XhoI) by Twist Bioscience or GenScript.

### Genome sequencing and proteomic analysis of Hydrogenophaga A37

*Hydrogenophaga* sp. A37 DNA was prepared and sequenced on MinION (R10.4 flow cells). Basecalling/demultiplexing used Dorado; reads were filtered with FiltLong and assembled with Flye, polished with Racon and Medaka, and evaluated with CheckM. GTDB-Tk and NCBI PGAP v6.10 were used for taxonomic assignment and annotation. (Assembly accession and metrics are reported in the SI.) For proteomic analysis, *Hydrogenophaga* sp. A37 was grown in 1/10 TSB; cells harvested in biological quadruplicate at OD600 ≈0.6. Proteins were extracted, trypsin digested in S-Trap, and analyzed by nano-LC-MS/MS on a Q-Exactive Orbitrap. Raw files were processed with FragPipe/MSFragger/IonQuant/Philosopher; searches used the new A37 proteome plus contaminants and decoys. Proteins were reported at 1% FDR and considered present if detected in at least 3 of 4 biological replicates.

### Screening of enzyme candidates and small-scale expression

Cloned constructs were transformed into *E. coli* BL21-Star (DE3). Expression screening used ZYP-5052 autoinduction medium supplemented with FeCl_3_ and 5-aminolevulinic acid (5-ALA); cultures were grown in deep-well plates overnight at 30 °C, lysed (BugBuster + lysozyme or sonication), and His-purified on Ni-IMAC magnetic beads. The activity screen used ABTS with added H_2_O_2_ (final ABTS 0.8 mM, H_2_O_2_ 1 mM) and monitoring at 418 nm.

### Production and purification of BUPOs and *Hsp*UPO

Selected BUPOs (*Age*BUPO, *Kal*BUPO, *Hyd*BUPO) were produced at liter scale in *E. coli* (ZYP-5052 autoinduction medium with FeCl_3_ and 5-ALA) and purified by HisTrap Ni-affinity and size-exclusion chromatography; *Kal*BUPO (pelB) was also recovered from culture medium when lysis/secretion occurred. *Hsp*UPO was expressed in *K. phaffii* (alpha factor signal peptide) and purified by ion exchange or Strep-tag affinity as described in SI. Heme-loaded protein concentrations were estimated by A_420_, and iron and magnesium quantification was performed by ICP-MS.

### Activity assays and analytics

Peroxidase/peroxygenase assays included ABTS, 2,6-DMP, indole, and naphthalene spectrophotometric assays (assay parameters and concentrations in SI). 3-phenyl-1-propanol (3PP) oxidations were followed by reversed phase HPLC (UV detection) with catalase quench for time courses. Reactions with palmitic acid and 16-hydroxypalmitic acid were analyzed by LC-MS (negative mode); products were identified using standards when available using extracted-ion chromatograms (m/z values and quantitation details in SI). H_2_O_2_-dependent inactivation of *Kal*BUPO was assayed by preincubation with 1 mM H_2_O_2_ prior to naphthalene activity measurement.

## Acknowledgments

This research was funded by the Research Council of Norway (grant 326975 to ELT, AAS, VGHE, SLLR, RMS and GV-K), a strategic PhD fellowship provided by NMBU and by NMBU’s SmartPlast sustainability arena (grant 1211130202D).

The authors thank Prof. Åsa Frostegård for providing the *Hydrogenophaga* sp. A37 strain and Marcus Torres Hansen for his contribution in cultivating and sampling of the bacterium. The authors also thank the National network of Advanced Proteomics Infrastructure (NAPI) supported by the Research Council of Norway (grant 295910) for access to proteomics infrastructure.

## Data availability

The Whole Genome Shotgun project has been deposited at DDBJ/ENA/GenBank under the accession JBRKPL000000000. The version described in this paper is version JBRKPL010000000. Nanopore sequencing raw data is available as part of NCBI BioProject PRJNA1337102. The mass spectrometry proteomics data has been deposited to the ProteomeXchange Consortium via the PRIDE (Proteomics Identification Database) partner repository with the data set identifier PXD069273.

## Author Contributions

E.L.T. designed and performed experiments, analyzed the data, wrote the first draft, and contributed to manuscript editing. A.A.S., N.S.E., M.B.R., and R.M.S. designed and performed experiments, analyzed the data, contributed to data interpretation, and manuscript editing. S.L.L.R., analyzed the data, contributed to data interpretation, and manuscript editing. M.S. and V.G.H.E. contributed to data interpretation, and manuscript editing. G.V.-K. supervised the project, conceptualized the work, contributed to data interpretation, and manuscript editing.

## Competing Interest Statement

The authors declare no competing interests.

## Notes

### Competing Interest Statement

The authors have declared no competing interest.

