## Supplementary Information for "Discovery and characterization of bacterial unspecific peroxygenase-like heme-thiolate enzymes"

**This PDF file includes:**

- Supporting materials and methods
- Figures S1 to S13
- Tables S1 to S2
- Legends for Datasets S1 to S2
- SI References

**Other supporting materials for this manuscript include the following:**

- Datasets S1 to S2

### Supporting Materials and Methods

**Structure-based search with FoldSeek and identification of the BUPO dataset.** Two short-type fungal UPOs, *Mro*UPO (PDB 7zbp) and *Hsp*UPO (PDB 7o1r), and one family II ("long") member, *Aae*UPO's evolved mutant PaDa-I (PDB 5OXU), were used as templates to search for bacterial homologs using FoldSeek (1). The search was performed against the AlphaFold database AFDB50, using 3Di/AA mode (local alignment using the 3Di alphabet + BLOSUM62 as described in (1)) and a taxonomic filter restricting the results to bacteria (eubacteria). Only hits with a probability greater than 0.5 and sequences longer than 150 amino acids were considered as positive hits. The resulting amino acid sequences (19 proteins in total) were searched in UniProt BLAST individually to expand the dataset; identified proteins with E values lower than $10^{-18}$ and longer than 150 AA were added to the original set, and duplicate sequences were removed using Cd-hit (2) with 100% identity cutoff. SignalP 6.0 (3) was used to predict the presence of secretion signals and InterPro Scan (4) was used to annotate protein domains.

To reduce the redundancy of the dataset prior to further analyses of the sequences, Cd-hit was used again, this time with an identity cutoff of 90%, resulting in a reduced set of 91 sequences.

**Phylogenetic analysis.** To explore the relatedness of the putative bacterial UPOs and fungal UPOs, it was necessary to remove the additional domains of multi-domain proteins; thus, the AlphaFold models of two bacterial proteins containing a C-terminal cytochrome C peroxidase domain were visually analyzed to determine the limit of the UPO-like domain. These two sequences were manually truncated and aligned with the reduced dataset of bacterial proteins Clustal Omega (5). This initial multiple sequence alignment (MSA) was truncated at the limit of the UPO-like domain and a new MSA was made, now including a set of 25 previously studied fungal UPOs (**Table S1**), using the online MAFFT server (6).

The MSA was used to calculate the pairwise sequence identity percentage using the following equation:

$$\%\ \text{identity} = \frac{\text{number of identical non-gap residue pairs}}{\text{length of the shortest sequence in the pair}} \times 100$$

Key regions of the alignment were used to make sequence logos using the online WebLogo 3 server (7). Before phylogenetic analysis, the 6 most divergent bacterial sequences were removed from the alignment and then, a neighbor-joining phylogenetic tree was built in the MAFFT server (6). with the alignment containing a total of 110 sequences, 85 bacterial and 25 fungal, using the JTT substitution model and 100 bootstrap resampling rounds. The tree was annotated with the online tool iTol (8) coupling it with the output from InterPro Scan and SignalP described above.

By analyzing the phylogenetic tree and the predicted structures, 13 putative bacterial UPOs were selected for expression in *E. coli*, aiming to cover sufficient diversity to increase the chances of successful expression.

**Structure prediction and analysis.** Structure predictions of the 85 BUPOs shown in the phylogenetic tree were obtained using the AlphaFold3 server (9), including a heme b cofactor and a $Mg^{2+}$ ion as ligands. The predicted structures were aligned to the crystal structure of *Hsp*UPO (PDB 7o1r) using the TM-align algorithm in the Python package tmtools, version 0.2.0 (10–12), from which the TM-scores relative to *Hsp*UPO were obtained.

**Materials and expression vectors.** All chemicals were purchased from Sigma-Aldrich (St. Louis, MO, USA), unless otherwise stated. Palmitic acid was purchased from Fluka Chemie (Buchs, Switzerland), and 5-aminolevulinic acid was purchased from Chem-Impex (Wood Dale, IL, USA). The *Hydrogenophaga sp.* A37 isolate (13) was kindly provided by Prof. Åsa Frostegård at the Norwegian University of Life Sciences.

Genes encoding putative bacterial UPOs were codon-optimized for expression in *E. coli*, using the GenSmart codon optimization tool (https://www.genscript.com/tools/gensmart-codon-optimization), with added 6-His tags either at the N-terminus or the C-terminus, chosen by inspecting their AlphaFold models to avoid adding tags at termini that were either buried or close to the active site tunnel. For the proteins with a predicted signal peptide, three versions were designed: native (with native signal peptide), truncated (removed signal peptide), and with a pelB signal peptide (pelB-SP). In the truncated and pelB-SP variants of predicted lipoproteins, the lipidation site cysteine was also excluded. The amino acid sequences of the expressed proteins are shown in **Table S2**. The constructs were synthesized and cloned into the pET-29b(+) vector (*Nde*I/*Xho*I restriction sites) by Twist Bioscience (South San Francisco, CA, USA) or GenScript (Rijswijk, Netherlands) (detailed in **Table S2**).

**Resequencing of the *Hydrogenophaga* sp. A37's genome, assembly and annotation.** H*ydrogenophaga sp.* A37 was cultured in 1/10 diluted TSB medium (Tryptic Soy Broth, Merck, Darmstadt, Germany) for 48 h at 22 °C with shaking at 200 RPM. Genomic DNA was extracted using the DNeasy PowerSoil Pro kit (QIAGEN) following the manufacturer's instructions. DNA concentration was measured using a NanoDrop One spectrophotometer and a Qubit 3.0 fluorometer with the dsDNA High Sensitivity assay kit (Thermo Fisher Scientific). DNA quality was assessed by gel electrophoresis on a Bio-Rad Gel Doc EZ Imager. As this genome was sequenced with other samples, a sequencing library was prepared using the Native Barcoding kit SQK-NBD114.24 (Oxford Nanopore Technologies), following the manufacturer's protocols. The library was loaded onto two FLO-MIN114 R10.4 flow cells and sequenced for 48 h on a MinION device using MinKNOW v4.0.5. POD5 files were base-called and demultiplexed with Dorado v0.5.0 (https://github.com/nanoporetech/dorado) using the super-accurate model (dna_r10.4.1_e8.2_400bps_sup@v4.3.0). Raw reads were quality-filtered with FiltLong v0.2.1 (https://github.com/rrwick/Filtlong) using the parameters --min_length 3000, --keep_percent 90 and assembled using Flye v2.9.2 (--nano_hq, --min-overlap 1500, default settings) (14). Contigs were initially polished with two consecutive rounds using Medaka v2.0.1 (-m r1041_e82_400bps_sup_v5.0.0; https://github.com/nanoporetech/medaka), followed by mapping raw reads to the consensus sequence generated by Medaka using minimap2 v2.28-r1209 (15).

The alignment was polished using Racon v1.5.0 (-m 8 -x -6 -g -8 -w 500) (https://github.com/isovic/racon). Assembly completeness was evaluated with CheckM (16). GTDB-Tk with GTDB release 220 was used to determine the taxonomic assignment (17). Functional annotations were obtained using the NCBI Prokaryotic Genome Annotation Pipeline (PGAP) v6.10 (18).

**Proteomic analysis of *Hydrogenophaga sp.* A37**. H*ydrogenophaga sp.* A37 was grown in quadruplicate in 1/10 TSB for 48 h at 22 °C with shaking at 200 RPM. Samples (5 ml) were harvested at the mid-exponential growth phase ($OD_{600}$ 0.6). Cells were separated from the supernatant by centrifugation (6,000 × g, 20 min, 4 °C), resuspended in 50 mM Tris-HCl pH 7.5, 100 mM NaCl, 0.1% (v/v) Triton X-100, 1 mM dithiothreitol (DTT) and disrupted by bead-beating (three 60 s cycles) with a FastPrep24 (MP Biomedicals, CA) at 6.5 m $s^{-1}$. After removal of cell debris by centrifugation (16,000 × g, 10 min, 4 °C), the proteins in the supernatant were precipitated by adding 50% ice-cold TCA with a 1:4 ratio and incubated at 4 °C overnight. Precipitated proteins were collected by centrifugation (15,000 × g, 15 min, 4 °C) and washed by adding 300 µL ice-cold wash buffer and centrifugation (15,000 × g, 15 min, 4 °C). The wash buffer was decanted before air-drying proteins. Proteins were resuspended in 46 µL lysis buffer (5% SDS and 50 mM triethylammonium bicarbonate pH 8.5). Protein digestion was performed using S-Trap™ Mini Columns (Protifi, Fairport, NY, USA) according to the manufacturer's instructions, with 20 mM DTT for reduction and 40 mM iodoacetamide for alkylation. The peptides were dried in a speed-vac and then dissolved in 1 % (v/v) formic acid. Samples were injected into an Ultimate 3000 nano ultra-high performance liquid chromatography system (Dionex, Sunnyvale CA, USA) connected to a Q-Exactive quadrupole-orbitrap mass spectrometer (Thermo Scientific, Bremen, Germany) equipped with a nano-electrospray ion source. Chromatographic separation was performed on a nanoViper Acclaim PepMap 100 C18 column (3 µm, 100 Å, 50 cm; Dionex, Sunnyvale, CA, USA) at a flow rate of 300 nL $min^{-1}$. Peptides were eluted using a gradient from 12 % to 43 % solvent B over 93 min, followed by an increase to 90% B in 6 min (solvent A: 0.1 % formic acid in water; solvent B: 80 % acetonitrile, 0.08% formic acid). The mass spectrometer operated in data-dependent mode, alternating between Orbitrap MS (R = 70,000) and HCD Orbitrap MS/MS (R = 35,000). AGC target was set to 1,000,000 charges with a maximum injection time of 128 ms. The 10 most intense ions were selected for fragmentation per cycle, with a 20 s dynamic exclusion for previously fragmented precursors.

MS raw files were processed using the FragPipe v21.1 (https://fragpipe.nesvilab.org/), with MSFragger v4.0 (19), IonQuant v1.10.12 (20), Philosopher v5.1.0 (21) for protein identification and label-free quantification (LFQ). MS and MS/MS spectra were searched against the complete *Hydrogenophaga sp.* A37 proteome (5,226 proteins, derived from the newly assembled genome), supplemented with common contaminants (e.g., keratins, trypsin, and bovine serum albumin) and a decoy database of reversed sequences to estimate false discovery rates (FDRs). The final protein library comprised 10,688 proteins. Trypsin was set as the proteolytic enzyme, with one missed cleavage allowed. Carbamidomethylation of cysteines was set as a fixed modification while variable modifications included methionine oxidation and pyro-glutamate formation at N-terminal glutamines. Protein identifications were filtered to achieve a 1% FDR. A protein was considered 'present' if detected in at least three of the four biological replicates.

**Screening enzyme candidates for activity**. *E. coli* BL21 star (DE3) competent cells (Thermo Fisher Scientific, Waltham, USA) were transformed with the plasmids and transformants were selected on lysogeny broth (LB) agar plates containing 50 µg mL$^{-1}$ kanamycin and 5 mg mL$^{-1}$ glucose to minimize basal expression of T7 polymerase. Liquid cultures were prepared in sterile 12-well plates with 1.5 mL LB with 50 µg mL$^{-1}$ kanamycin and 5 mg mL$^{-1}$ glucose per well. Each well was inoculated with a single colony, and the plates were incubated overnight in thermomixers (Eppendorf, Hamburg, Germany) at 37 °C and 300 RPM. The resulting cultures were mixed 7:3 with sterile 86 % (w/w) glycerol in a polypropylene 96-well plate, sealed and stored at -80 °C until further use.

Expression tests were carried out using sterile 1.3-mL 96-deep-well plates and ZYP-5052 auto-induction medium (22), consisting of 5 mg mL$^{-1}$ yeast extract, 10 mg mL$^{-1}$ tryptone, 50 mM $Na_2HPO_4$, 50 mM $NaH_2PO_4$, 25 mM $NH_4SO_4$, 5 mg mL$^{-1}$ glycerol, 0.5 mg mL$^{-1}$ glucose, 2 mg mL$^{-1}$ lactose and 2 mM $MgSO_4$, with 200 µg mL$^{-1}$ kanamycin for selection, and 50 µM $FeCl_3$ and 500 µM 5-aminolevulinic acid to promote heme biosynthesis. The glycerol stocks were thawed and 3 µL of each clone was used to inoculate 600 µL of medium. For each clone two wells were inoculated and two wells inoculated with non-transformed *E. coli* BL21 star (DE3) were included, using the same medium but without kanamycin. The plate was sealed with a breathable sterile rayon film (Nunc, ThermoFisher; Rochester, NY, USA) and incubated at 30 °C for 24 hours in a thermomixer set to 800 RPM. The cultures were then transferred into 1.5 mL microcentrifuge tubes. The duplicate samples corresponding to the same experimental conditions (i.e., the same enzyme variants or the same control cultures) were pooled together resulting in 1.2 mL final volume. Next, the cells were pelleted by centrifugation (10 minutes at 14,000 × g) and the pellets were stored at -20 °C until further use.

Cell lysis was performed by resuspending the cells in 250 µL of BugBuster protein extraction reagent (Merck, Darmstadt, Germany) containing 0.4 mg mL$^{-1}$ lysozyme (Roche Diagnostics, Mannheim, Germany). The cell suspensions were incubated in a thermomixer at 25 °C and 300 RPM for 30 minutes, followed by centrifugation (20 minutes at 20,000 × g). Proteins were purified from the supernatants using Pierce High-Capacity Ni-IMAC magnetic beads (Thermo Fisher Scientific; Waltham, MA, USA). 20 µL of a 25% (w/w) beads suspension was loaded into 1.5 mL microcentrifuge tubes and equilibrated with 500 µL of 50 mM sodium phosphate buffer, pH 7.4 containing 5 mM imidazole and 300 mM NaCl. The tubes were placed on a magnetic rack and the supernatants were replaced by 750 µL of the same equilibration buffer. 250 µL of clarified lysates was added into each tube followed by incubation for 10 minutes at room temperature in a rotary shaker (Multi RS-60; BioSan, Riga, Latvia) set to 25 RPM. The supernatants were removed using the magnetic rack, and the beads were washed two times with 500 µL of 50 mM sodium phosphate buffer, pH 7.4 containing 10 mM imidazole and 300 mM NaCl. The proteins were eluted with 200 µL of 50 mM sodium phosphate buffer, pH 7.4 supplied with 500 mM imidazole and 300 mM NaCl.

To probe enzymatic activity, 10 µL of the resulting protein samples was transferred into 96-well microtiter plates and mixed with 80 µL of a 1 mM 2,2'-azino-bis(3-ethylbenzothiazoline-6-sulfonic acid (ABTS) solution in 50 mM sodium phosphate buffer, pH 6.0. The reactions were started by adding 10 µL of a 10 mM $H_2O_2$ solution in Milli-Q water followed by mixing for 30 seconds at 600 RPM using a Varioskan LUX plate reader (Thermo Fisher Scientific, Waltham, MA, USA). The final concentration of ABTS and $H_2O_2$ in the reaction mixtures was 0.8 mM and 1 mM, respectively. The experiments were carried out for 1 hour at 25 °C. Product formation (i.e., the generation of ABTS radicals) was followed by measuring the optical absorbance at 418 nm.

**Production and purification of BUPOs.** The glycerol stocks of the strains expressing *Age*BUPO (native variant), *Kal*BUPO (pelB-SP variant), and *Hyd*BUPO (native variant) were used to start 20-mL pre-cultures in 100-mL flasks containing LB medium supplemented with 50 µg $mL^{-1}$ kanamycin and 5 mg $mL^{-1}$ glucose. The pre-cultures were incubated overnight at 37 °C and 200 RPM in an Ecotron incubator (Infors HT, Bottmingen, Switzerland). 10 mL of the overnight pre-cultures was used to inoculate shake flasks containing 1 L of ZYP-5052 auto-induction medium, supplemented with 500 µM δ-aminolevulinic acid, followed by incubation at 200 RPM and 30°C for 24 h (*Age*BUPO and *Hyd*BUPO), or 16 °C for 96 h (*Kal*BUPO) in an Ecotron incubator. Afterwards, the cultures were harvested by centrifuging at 8,000 × g for 10 min. The pellets from bacteria expressing *Age*BUPO and *Hyd*BUPO were resuspended in metal affinity chromatography binding buffer (20 mM sodium phosphate, 0.5 M NaCl, 40 mM imidazole, pH 7.4) and lysed by sonicating with a VibraCell ultrasonic disintegrator equipped with a micro tip probe (Sonics, Newtown, CT, USA), for 10 min at 28 % amplitude, pulsing 5 s on and 5 s off. The lysates were clarified by centrifugation for 10 min at 20,000 × g at 4 °C. The clarified lysates were manually loaded with a syringe onto a 5-mL HisTrap FF Crude column (Cytiva, Marlborough, MA, USA), pre-equilibrated with binding buffer. The columns were then washed with 10 column volumes of binding buffer, after which the proteins were eluted with 5 column volumes of elution buffer (20 mM sodium phosphate, 0.5 M NaCl, 500 mM imidazole, pH 7.4).

Expression of *Kal*BUPO with the pelB signal peptide seemingly resulted in cell lysis during growth, evidenced by the dark red color of the clarified medium, and thus the enzyme was purified directly from the clarified medium by directly loading it (1 L) onto a pre-equilibrated 5-mL HisTrap FF Crude column (Cytiva) using a BioLogic LP System (Bio-Rad, Hercules, CA, USA). The column was then washed with 10 column volumes of binding buffer and the UPO was eluted with 10 column volumes of elution buffer.

The proteins were concentrated to < 2 mL using Amicon® Ultra-15 centrifugal filters (Merck KGaA, Darmstadt, Germany) with a molecular weight cutoff of 10 kDa and filtered through 0.22 µm filters to remove potential protein precipitates. Each protein was then further purified by size-exclusion chromatography using a ProteoSEC Dynamic 16/60 3–70 HR column (Protein Ark, Sheffield, UK), operated at 1 mL $min^{-1}$ flow rate and equilibrated with 20 mM sodium phosphate buffer, pH 7.0, containing 200 mM NaCl. The fractions showing absorbance at 420 nm were pooled and concentrated as described above, and the heme-containing protein concentration was estimated by measuring the absorbance at 420 nm (reflecting the presence of UPO bound heme) using the extinction coefficient of *Mro*UPO ($\varepsilon_{420}$ = 115 $mM^{-1}$ $cm^{-1}$) (23), assuming similar extinction coefficients for the BUPOs. The correct extinction coefficients were later determined using the iron quantification through ICP-MS, confirming that they are close to that of *Mro*UPO (see below in section "**Quantification of iron and magnesium with ICP-MS"**).

**Production and purification of *Hsp*UPO.** The gene encoding the fungal UPO from *Hypoxylon sp., Hsp*UPO (UniProt accession A0A1Y2TH07) was codon-optimized for expression in *Komagataella phaffii* and synthesized by Twist Biosciences (South San Francisco, CA, USA), replacing the native signal peptide sequence by that of the alpha factor signal peptide, and including 30 bp overhangs on each end, homologous to the expression vector pBSY3Z (Bisy GmbH, Hofstaetten a. d. Raab, Austria) for Gibson assembly cloning. The vector was linearized by PCR using the primers 5'-TTTAATTGTAAGTCTTGACTAGAGCAAGTG-3' and 5'-GCGGCCGCTCAAGAGGAT-3' and the Q5 High-Fidelity polymerase (New England Biolabs, USA), treated with the restriction enzyme *Dpn*I (New England Biolabs, USA), and purified with the DNA Clean-up and Concentration kit (ZymoResearch, USA). The *Hsp*UPO gene fragment was then assembled into the expression vector by using a NEBuilder HiFi DNA Assembly kit (New England Biolabs, USA) according to the manufacturer's instructions. The assembled DNA was introduced into *E. coli* TOP10 cells (Invitrogen, Thermo Fisher Scientific, USA) and transformants were selected by plating on LB agar plates containing 25 µg $mL^{-1}$ of Zeocin (Gibco, Thermo Fisher Scientific, USA). Single colonies were cultured overnight in 5 mL LB containing 25 µg $mL^{-1}$ of Zeocin for plasmid propagation, after which plasmid DNA was purified with the EZNA Plasmid DNA Mini Kit I (Omega Bio-Tek, USA) and then sequenced by Eurofins Genomics (Ebersberg, Germany) using Sanger sequencing. A version of the gene with an added affinity tag StrepTagII (WSHPQFEK) at the N-terminus of the UPO, downstream of the signal peptide, was produced by amplifying the sequence-verified plasmid the primers 5'-TTTTTCGAACTGCGGGTGAGACCA-AGCTTCGGCCTCTCTCTTCTCG-3' and 5'-CTTGGTCTCACCCGCAGTTCGAAAAAGCTCCAT-CTCCATCTTCTGGTTG-3', following the same procedure for cloning, selection and sequence verification as described above. Sequence-verified plasmids were then linearized with *Swa*I (New

England Biolabs, USA) and used to transform electrocompetent *K. phaffii* MutS BSYBG11 cells (Bisy GmbH, Austria). Successful transformants were selected on YPD agar plates containing 100 µg mL$^{-1}$ Zeocin. Single colonies were used to inoculate 5-mL cultures in BMD1 medium (200 mM potassium phosphate buffer pH 6.0, 13.4 g L$^{-1}$ yeast nitrogen base, 0.4 mg L$^{-1}$ biotin, 10 g L$^{-1}$ glucose), which were incubated at 30 °C and 200 RPM for 66 hours. After that, 0.5 mL of BMM10 medium (200 mM potassium phosphate buffer pH 6.0, 13.4 g L$^{-1}$ yeast nitrogen base, 0.4 mg L$^{-1}$ biotin, 5% (v/v) methanol) was added for induction, with subsequent additions of 50 µL of pure methanol at 74, 90, and 98 h. Cultures were harvested at 114 h by centrifugation (5,000 × g, 10 min, at 4 °C). The supernatants were used carry out an activity assay with 2,6-DMP (2,6-dimethoxyphenol, Merck, Germany), as described below in the section "**Peroxidase colorimetric assay with 2,6-DMP**" and the clone with highest activity was used to produce the enzyme at larger scale. Then, *Hsp*UPO and *Hsp*UPO-StrepTag**II** were produced in 2-L baffled Erlenmeyer flasks containing 500 mL of BMD1 medium. The cultures were incubated at 30 °C and 200 RPM for 66 hours, after which 50 mL of BMM10 medium was added for induction, with subsequent additions of 2.5 mL of pure methanol every 12 hours. At 160 hours, the culture medium was harvested by centrifugation at 8,000 × g and 4 °C for 20 minutes. The cell pellet was discarded, and the supernatant was filtered through a 0.22 µm filter (SteriTop, Merck, Rahway, NJ, USA) and concentrated 10-fold using a Tangential Flow Filtration (TFF) cassette (Vivaflow, Sartorius, Göttingen, Germany).

*Hsp*UPO (with no affinity tag) was purified from the concentrated culture supernatant using ion exchange chromatography (HiTrap Capto Q, 5 mL; Cytiva, Marlborough, MA, USA), with the column pre-equilibrated in 20 mM Tris-HCl buffer, pH 8.0 and proteins eluted with a 0–0.5 M NaCl gradient over 40 column volumes. *Hsp*UPO-StrepTag**II** was purified by loading the concentrated culture supernatant onto a 5-mL StrepTrap XT affinity column (Cytiva, Marlborough, MA, USA), pre-equilibrated in 100 mM Tris-HCl, 150 mM NaCl, 1 mM EDTA, pH 8, and proteins were eluted using 3 column volumes of a buffer with the same composition but containing 50 mM biotin. Fractions with the highest absorbance at 420 nm were pooled, concentrated and buffer-exchanged into 20 mM sodium phosphate, pH 7.0, using an Amicon® Ultra-15 centrifugal filter (Merck KGaA, Darmstadt, Germany) with a molecular weight cutoff of 10 kDa. The protein concentration was estimated by measuring absorbance at 420 nm, using the extinction coefficient of *Mro*UPO (115 mM$^{-1}$ cm$^{-1}$ (23)), assuming a similar value for *Hsp*UPO. The correct extinction coefficient was later determined using the iron quantification through ICP-MS (see below, in section "**Quantification of iron and magnesium with ICP-MS"**).

**UV-Vis spectra.** The UV-Vis spectra of *Age*BUPO, *Kal*BUPO and *Hyd*BUPO were recorded using a Cary 60 UV-Vis spectrophotometer (Agilent, Santa Clara, CA, USA). The enzyme preparations were diluted to 5-10 µM (based on the absorbance at 420 nm) in 20 mM sodium phosphate buffer, pH 7.0, and the resting state spectra were recorded in a quartz cuvette with 1 cm pathlength. Then, a few grains of sodium dithionite were added to the cuvette, and the sample was gently mixed by inversion before measuring the UV-Vis spectrum again, yielding the spectrum for the reduced enzyme.

**Quantification of iron and magnesium with ICP-MS.** *Age*BUPO, *Kal*BUPO, *Hyd*BUPO and *Hsp*UPO-StrepTagII were diluted in 20 mM sodium phosphate buffer pH 7.0 to 1-2 µM heme-containing protein using the concentration estimated with the absorbance at 420 nm as described above. Buffer alone was used to quantify the background Fe and Mg content, and deionized water was used as blank for the analysis. One milliliter of each sample (in triplicate) was weighed in 15-mL centrifuge tubes, and 0.75 mL of ultrapure concentrated $HNO_3$ was added. The samples were heated to 90 °C for 1 h and then diluted with deionized water to a final volume of 10 mL. The concentrations of Fe and Mg were quantified using a triple-quadrupole ICP-MS system (8900 ICP-QQQ, Agilent, Santa Clara, CA, USA) operated in He-KED mode at masses 56 and 24 amu for Fe and Mg, respectively, with indium as the internal standard. The limits of detection (3 × SD from blank samples, n=5) and quantification (10 × SD from the blank samples, n=5) were 0.0007 and 0.0023 mg $kg^{-1}$ for Fe, and 0.0007 and 0.0022 mg $kg^{-1}$ for Mg, respectively.

**Peroxidase colorimetric assay with ABTS.** The reactions were carried out in 96-well plates, in a total volume of 200 µL, containing 0.8 mM ABTS and 0.5 µM enzyme in 100 mM sodium citrate buffer, pH 5.5. The reference enzyme, *Hsp*UPO, was used at 0.05 µM. Control reactions with 0.5 µM hemin chloride or buffer instead of enzyme, were included. The experiment was started by dispensing 10 µL of 20 mM $H_2O_2$ (to a final concentration of 1 mM), or water, in control reactions, using a Varioskan Lux plate reader (Thermo Fisher Scientific), and product formation was monitored by measuring the absorbance at 418 nm for 15 minutes, using the built-in pathlength correction method. The concentration of oxidized ABTS was calculated using the molar extinction coefficient $\varepsilon_{418}$ = 36 $mM^{-1}$ $cm^{-1}$ (24).

**Peroxidase colorimetric assay with 2,6-DMP.** In the peroxidase reaction with 2,6-dimethoxyphenol (2,6-DMP), the formation of the dimerized product coerulignone was monitored following absorbance at 469 nm ($\varepsilon_{469}$ = 27.5 $mM^{-1}$ $cm^{-1}$ (24)). The reaction mixture contained 2 mM 2,6-DMP, 2 mM $H_2O_2$, 100 mM potassium phosphate buffer, pH 6.0, and 0.5 µM bacterial enzyme or 0.05 µM *Hsp*UPO. Control reactions with 5 µM hemin chloride, 5 µM $FeCl_3$ or buffer instead of enzyme were included. 20 µL of 10 x enzyme solution was added in a 96-well plate and the experiment was started by dispensing 180 µL of a reaction mix containing the rest of the components using a Varioskan Lux plate reader (Thermo Fisher Scientific; Rochester, NY, USA). The absorbance at 670 nm was monitored for 15 minutes, using the built-in pathlength correction method.

**Peroxygenase colorimetric assay with indole.** The peroxygenase reaction with indole, leading to the formation of indigo, was monitored by measuring absorbance at 670 nm ($\varepsilon_{670}$ = 4.8 $mM^{-1}$ $cm^{-1}$ (24). The reaction mixture contained 2 mM indole (diluted 10 times from a stock prepared in 50 % (v/v) acetonitrile, leading to a final acetonitrile content of 5 %), 2 mM $H_2O_2$, 100 mM potassium phosphate buffer, pH 7.0, and 5 µM bacterial enzyme or 0.5 µM *Hsp*UPO. Control reactions with 5 µM hemin chloride, 5 µM $FeCl_3$ or buffer instead of enzyme were included. 20 µL of a 10 x enzyme solution was added in a 96-well plate and the experiment was started by dispensing 180 µL of a reaction mix containing the rest of the components using a Varioskan Lux plate reader (Thermo Fisher Scientific; Rochester, NY, USA). The absorbance at 670 nm was monitored for 15 minutes, using the built-in pathlength correction method.

**Peroxygenase spectrophotometric assay with naphthalene.** In the peroxygenase reactions with naphthalene, the formation of 1-naphthol was monitored by measuring the absorbance at 324 nm. The reaction mixtures contained 1 mM naphthalene, 5 µM bacterial enzyme or 0.5 µM *Hsp*UPO, 100 mM sodium citrate buffer, pH 5.5, and 10 % acetone. Control reactions with 5 µM hemin chloride, 5 µM $FeCl_3$ or water instead of enzyme were included. The experiment was started by dispensing 10 µL of 20 mM $H_2O_2$ (to a final concentration of 1 mM), or buffer, using a Varioskan Lux plate reader (Thermo Fisher Scientific; Rochester, NY, USA), and the absorbance at 324 nm was monitored for 15 minutes. A standard curve of 1-naphthol (5-1000 µM) was used to calculate the concentration of product.

**Monitoring the enzymatic oxidation of 3-phenyl-1-propanol (3PP).** The oxidation of 3PP by BUPOs and *Hsp*UPO was assessed in 50 mM sodium phosphate buffer, pH 6.0, supplied with 1 mM substrate and 5 µM enzyme. The experiments were initiated by adding $H_2O_2$ to 1 mM final concentration and were carried out at 30 °C (using a thermomixer; Eppendorf, Hamburg, Germany) or at the room temperature. Reactions with hemin chloride substituting the enzyme and reactions with just $H_2O_2$ and the substrate were used as negative controls. For endpoint experiments, the reaction samples were transferred to HPLC vials and immediately analyzed as described below. For time-course experiments, aliquots were taken at various time points, and the reactions were quenched by adding bovine liver catalase (Sigma-Aldrich, St. Louis, MO, USA) to a final concentration of 250 u $mL^{-1}$. The reaction mixtures were analyzed by reverse phase chromatography using a Dionex UltiMate 3000 RSLC system (Thermo Fisher Scientific, Waltham, MA, USA) equipped with a Zorbax RR Eclipse Plus C18 column (2.1 × 150 mm, 3.5 µm) and a Zorbax RRHD Eclipse Plus C18 guard column (2.1 × 5 mm, 1.8 µm) produced by Agilent (Santa Clara, CA, USA). Separation was carried out using 10 mM ammonium acetate (buffer A) and 100% acetonitrile (buffer B) at 0.35 mL $min^{-1}$ flow rate. The column was equilibrated for 5 minutes (99 % buffer A, 1 % buffer B) and then a linear gradient of buffer B (1-25 %) was applied over 20 minutes at a column temperature of 30 °C. 3PP and its oxidized derivatives were detected by monitoring the UV absorbance at 260 nm.

**Oxidation of palmitic acid and hydroxy-palmitic acid.** Enzymatic reactions with palmitic acid and 16-hydroxypalmitic acid as substrates were carried out in 1.5-mL tubes, in a total volume of 100 µL, containing 5 µM of enzyme (except *Hsp*UPO, which was used at 0.5 µM) or hemin chloride,

0.1 mM substrate, 20 % (v/v) acetone, and 10 mM ammonium acetate, pH 6.0. The experiments were started by adding $H_2O_2$ to a final concentration of 1 mM followed by incubation at 30 °C and 1,000 RPM in a thermomixer (Eppendorf, Hamburg, Germany) for 1 h. Next, 200 µL of 90 % (v/v) acetonitrile containing 6 mM ammonium acetate was added, and the reactions were stirred for 5 min at 1,000 RPM in the thermomixer at 20 °C, to allow protein to precipitate and favor the dissolution of the analytes, followed by centrifuging for 5 min at 20,000 × g to remove protein precipitates. 60 µL of the supernatant was transferred to HPLC vials and the samples were analyzed by LC-CAD/MS using a Dionex UltiMate 3000 RSLC system (Thermo Fisher Scientific, Waltham, MA, USA) equipped with a Zorbax RR Eclipse Plus C18 column (2.1 × 150 mm, 3.5 µm) and a Zorbax RRHD Eclipse Plus C18 guard column (2.1 × 5 mm, 1.8 µm) produced by Agilent (Agilent, Santa Clara, CA, USA). The chromatography system was coupled, with a flow splitter 1:1, to a Corona Ultra charged aerosol detector (CAD) and a Velos Pro MS system (Thermo Fisher Scientific, Waltham, MA, USA). Chromatographic separation of the analytes was performed using a linear gradient from 50% to 99% eluent B (acetonitrile) over 2 minutes, followed by an isocratic hold at 99% B for 6 minutes, at a flow rate of 0.4 mL $min^{-1}$. Eluent A was 10 mM ammonium acetate in water. The MS system was operated in negative ionization mode with a scanning range of 500-1,000 m/z. The CAD nebulizer temperature was set to 30 °C. 16-hydroxyhexadecanoic acid, and hexadecanedioic acid were used as standards to identify elution peaks and quantify the corresponding compounds by integrating the peak areas in the extracted ion chromatograms of their respective $[M - H]^-$ m/z values (271.23 ± 0.2, and 285.21 ± 0.2). The amounts of keto and aldehyde products (isomeric) were estimated by integrating the peaks at m/z 269.21 ± 0.2 and calculating the concentrations using the standard curve of 16-hydroxypalmitic acid, assuming the same response factor, due to the unavailability of authentic standards.

**$H_2O_2$-dependent inactivation of *Kal*BUPO.** The inactivation of *Kal*BUPO was investigating by pre-incubating 20 µM *Kal*BUPO with 1 mM $H_2O_2$ for increasing times before measuring the residual activity on naphthalene. 33.3 µL of a 30 µM *Kal*BUPO solution was added in several wells of a 96-well plate, and a Varioskan LUX plate reader (Thermo Fisher Scientific, Waltham, MA, USA) was used to dispense 17 µL of 3 mM $H_2O_2$ to each enzyme-containing well (final volume 50.3 µL, 1 mM $H_2O_2$, 20 µM *Kal*BUPO) at 30, 20, 10, 5, 1 and 0 minutes before the addition of 150 µL of naphthalene reaction mix. The final composition of the 200 µL reaction mixture was 1 mM naphthalene, 10 % acetone, 100 mM citrate buffer, pH 5.5, and 1 mM freshly added $H_2O_2$ plus up to 0.25 mM $H_2O_2$ remaining from the pre-incubation, some of which might have been consumed by the enzyme. A control reaction was done without pre-added $H_2O_2$ (1 mM final $H_2O_2$) to account for the difference in $H_2O_2$ concentration at the start of the naphthalene oxidation reaction.

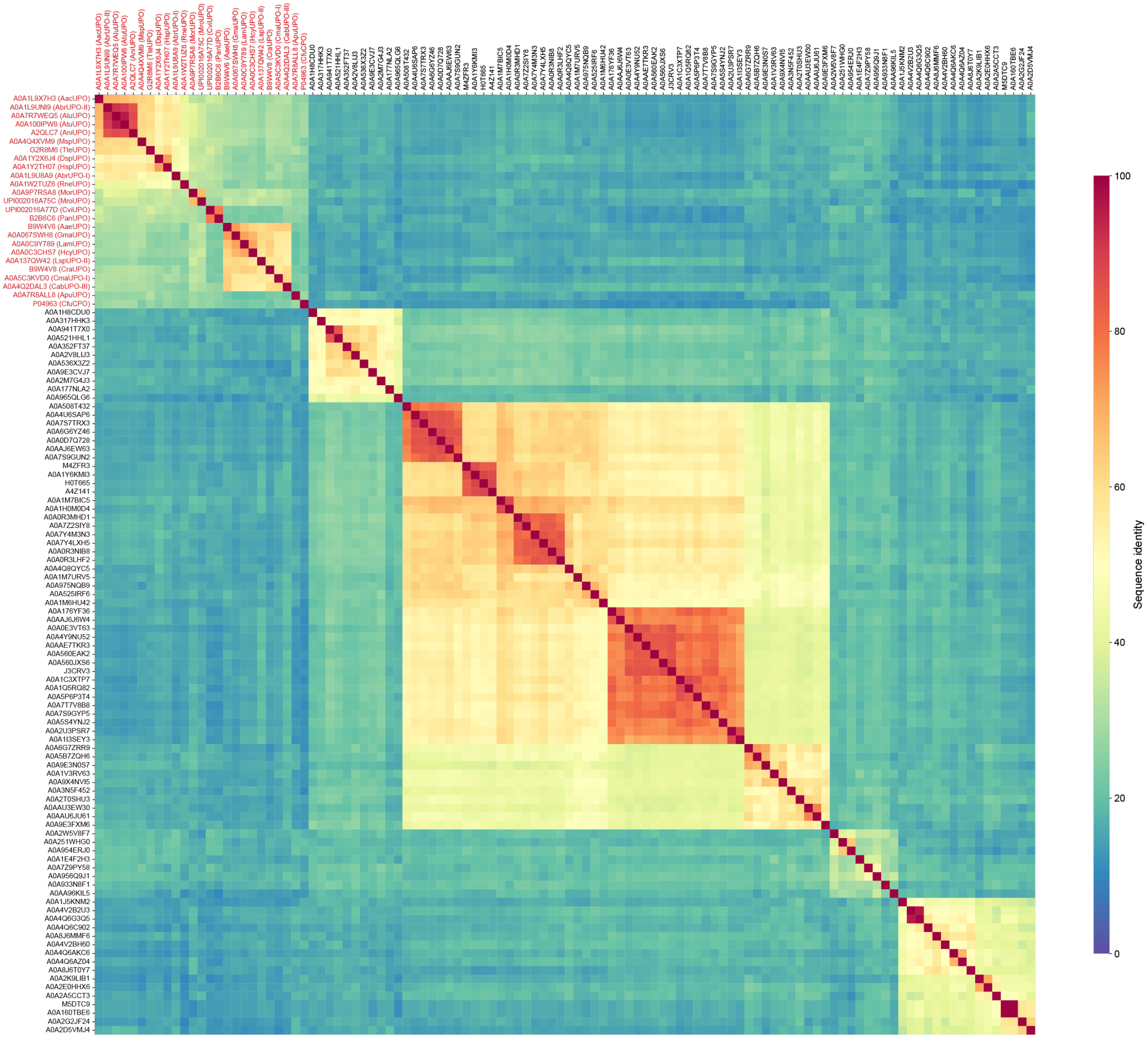


**Fig. S1. Sequence identity matrix of fungal UPOs and putative bacterial UPOs.** The sequence identity for each protein pair is color coded according to the scale shown next to the heatmap. Fungal protein identifiers are shown as red colored text. The matrix was calculated from the multiple sequence alignment described in Supplementary Methods, where additional C-terminal domains, if present, were removed from the BUPO sequences before the alignment was made. The percent sequence identity (SID) was calculated as %SID = IAP × $L^{-1}$ × 100, where IAP is the number of identical aligned non-gap residue pairs, and L is the length of the shortest protein of the pair.

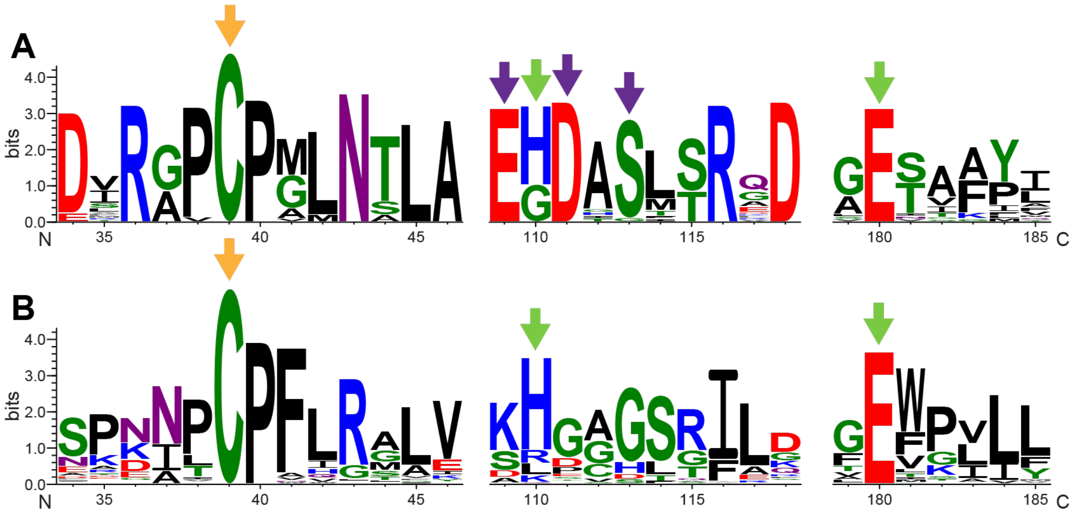


**Fig. S2. Sequence logos of conserved motifs in UPOs.** Panel **A** shows key conserved features in fungal UPOs; specific (putative) functions are indicated with arrows as follows: **orange**, heme-coordinating cysteine; **purple**, $Mg^{2+}$-coordinating residues; **lime green**, acid-base catalytic pair (position His110 is exclusive to short UPOs; this base is replaced by an arginine in long UPOs, located in a region that is not shown in this figure). Panel **B** shows the equivalent positions in putative bacterial UPOs, which have the heme-coordinating cysteine and the acid-base catalytic pair, but lack the $Mg^{2+}$-coordinating residues. The positions shown on the horizontal axis relate to the sequence of *Hsp*UPO, including its secretion signal peptide. All logos were generated using WebLogo 3 (7) from a multiple sequence alignment made with 25 fungal and 85 bacterial sequences (see Supplementary Methods for details).

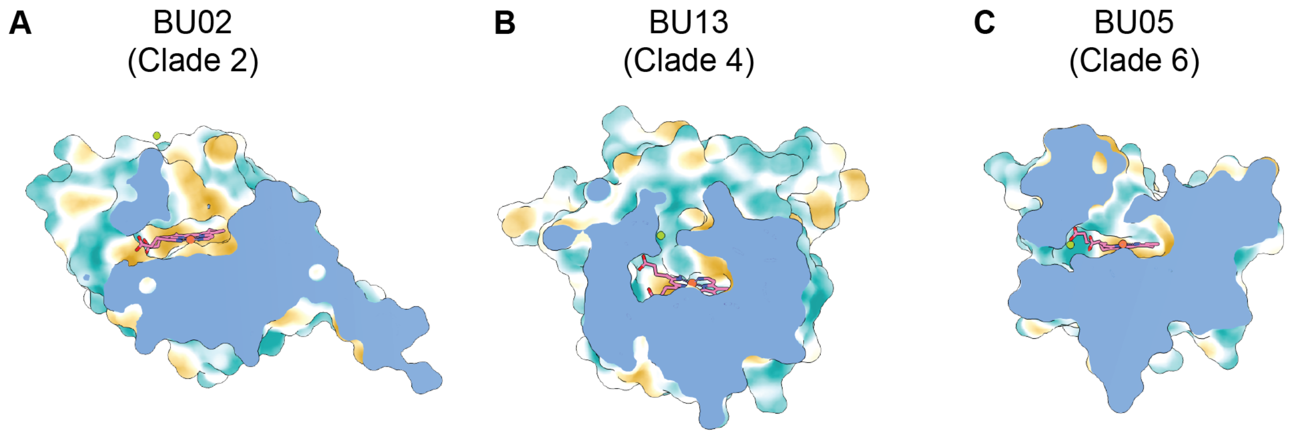


**Fig. S3. Active site access tunnels in BUPOs from clades 2, 4, and 6.** The tunnels leading to the heme cofactor are shown by protein structure cross-sections for representatives of the clades not shown in **Fig. 2**. Panel **A** shows BU02 (UniProt accession A0A4Q6C902), **B** shows BU13 (A0A2V8LIJ3), and **C** shows BU05 (A0A0R3MHD1). Note that BU13 (**B**), a representative of clade 4, does not feature a second entrance tunnel to the active site as the rest of the clades. The solvent excluded surface is shown in a lipophilicity color scale, from gold (lipophilic) to teal (hydrophilic). The models colored by pLDDT score are shown in **Fig. S4**.

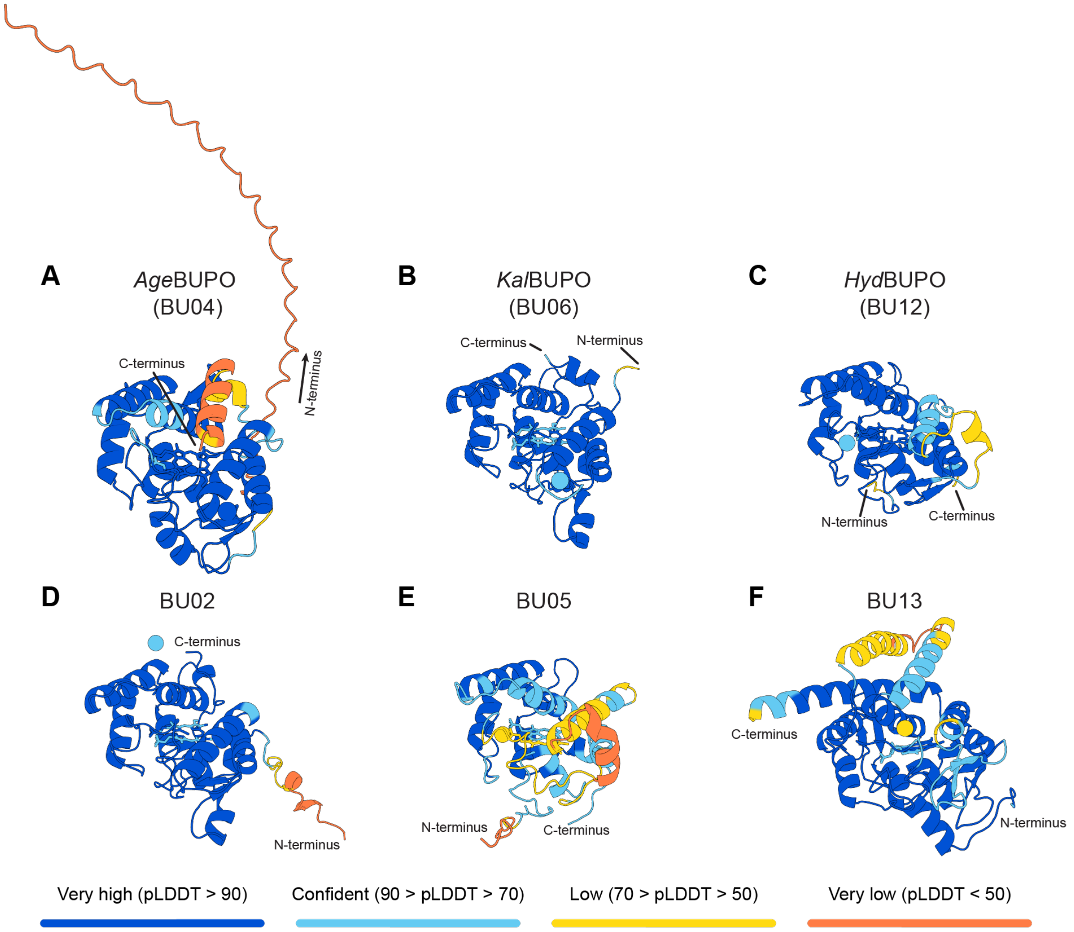


**Fig. S4. Structures of putative BUPOs predicted by AlphaFold 3.** The structure models used in **Figures 2** and **S3** are shown colored by pLDDT values along the peptide chain and ligands (heme b cofactor and $Mg^{2+}$ ion), using the Alphafold color scale. Panels **A-F** show the structures of *Age*BUPO (UniProt accession A0A2W5V8F7, clade 3), *Kal*BUPO (A0A2K9LIB1, clade 1), *Hyd*BUPO (A0A1V3RV63, clade 5), BU02 (A0A4Q6C902, clade 2), BU05 (A0A0R3MHD1, clade 6), and BU13 (A0A2V8LIJ3, clade 4), respectively.

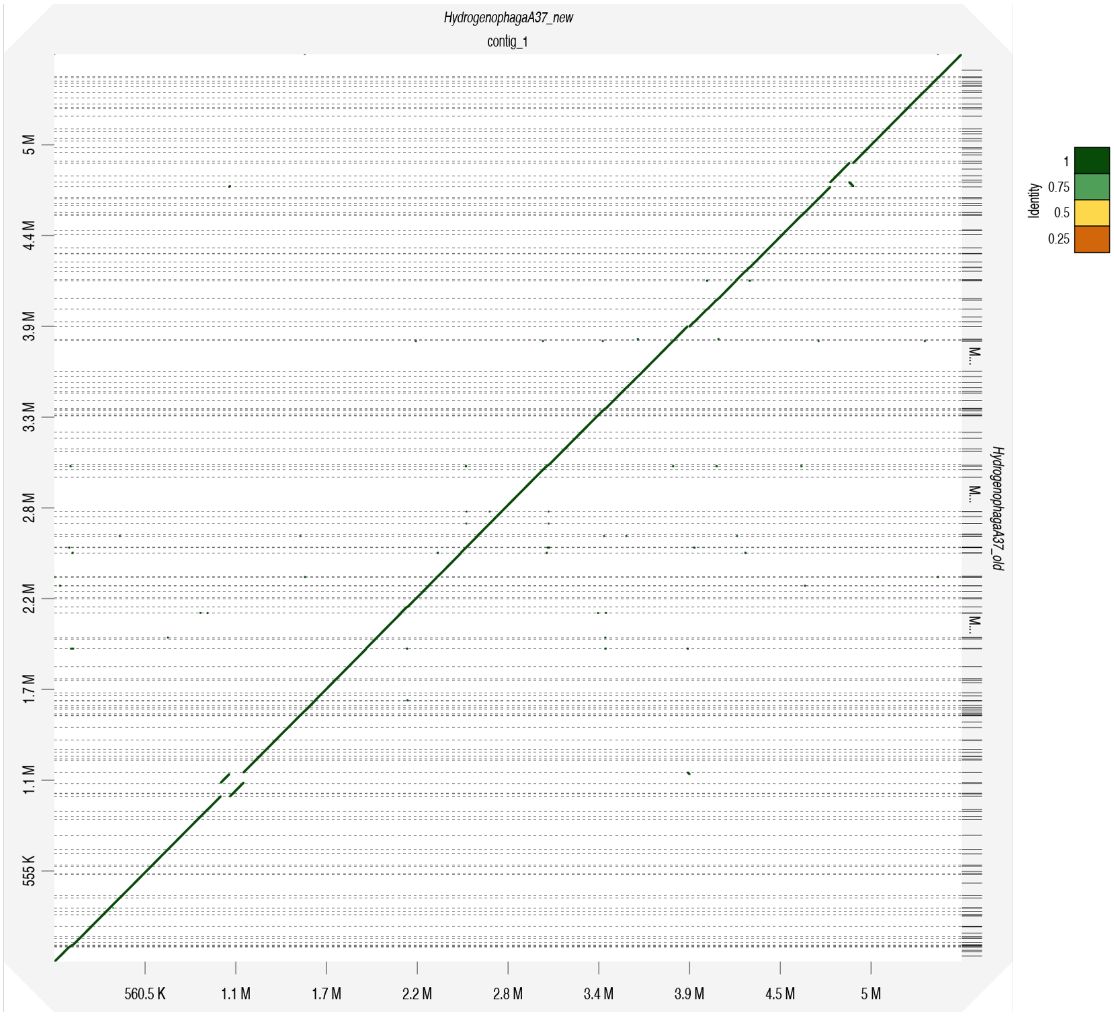


**Fig. S5. Alignment between the previous and new version of the genome of *Hydrogenophaga sp.* A37**. The new version of the genome (GCA_053010425.1) is shown on top, while the previous version (GCA_002001205.1) is shown on the right. The alignment was visualized with D-GENIES.

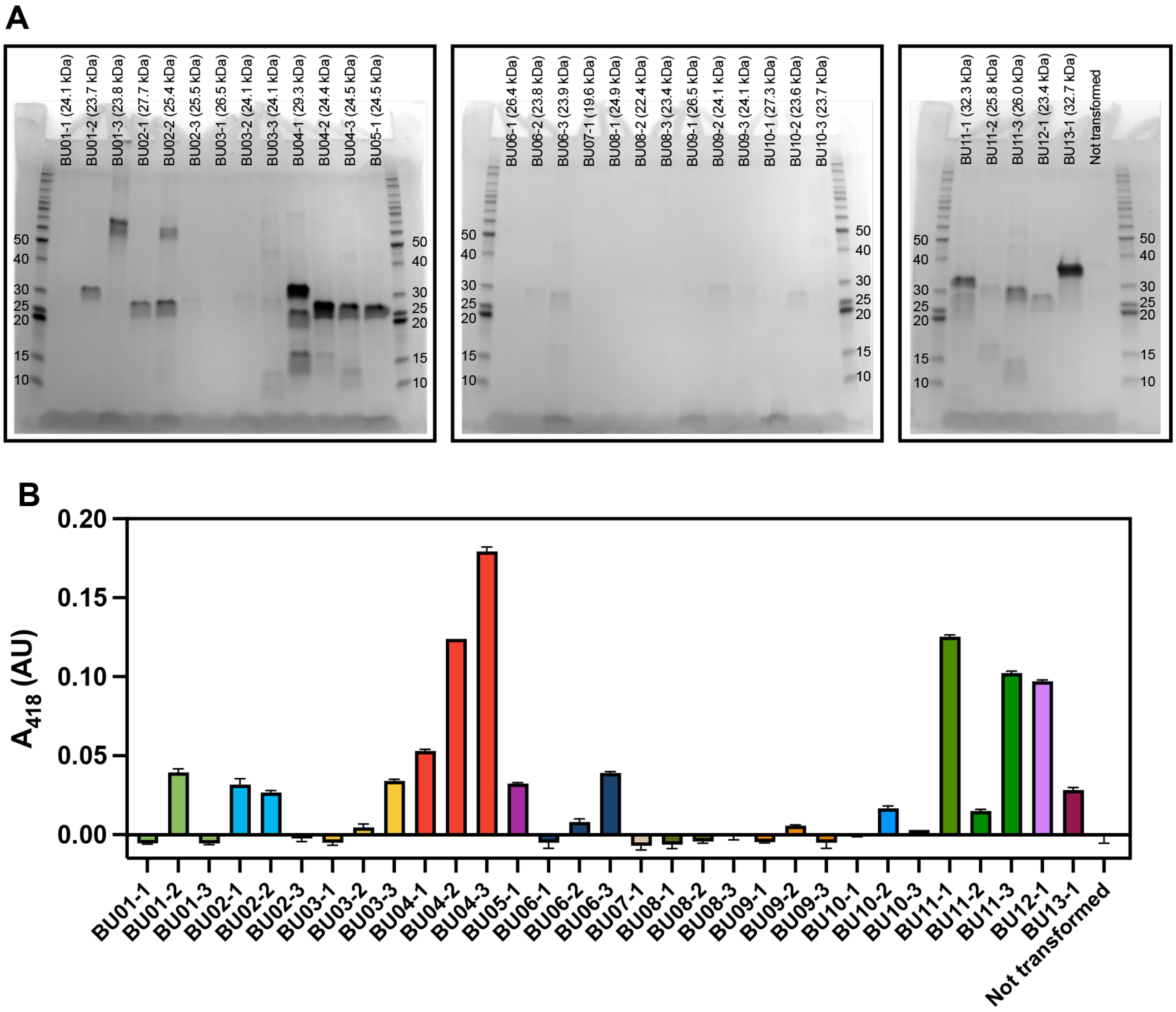


**Fig. S6. Expression and peroxidase activity of 13 putative bacterial UPOs.** Panel **A** shows SDS-PAGE analysis of partially purified putative BUPOs produced recombinantly in *E. coli* (see **Table S2** for details of the plasmids used). The gel was stained with Imperial Protein Stain (Thermo Fisher Scientific). The sizes of the protein MW standard (Benchmark protein ladder, Thermo Fisher Scientific) are indicated in kDa next to the corresponding bands, and the expected size of each protein is indicated in parentheses on top of each lane. Panel **B** shows the peroxidase activity in the partially purified protein solutions. Here, 10 μL of each protein solution was used in a peroxidase activity assay with 0.8 mM ABTS and 1 mM $H_2O_2$ in 50 mM sodium phosphate buffer, pH 6.0, in a final volume of 100 μL. The reactions were incubated for 1h at room temperature and product formation was determined by measuring absorbance at 418 nm. The absorbance values measured immediately after starting the reactions were subtracted from the end point measurements to account for background signals.

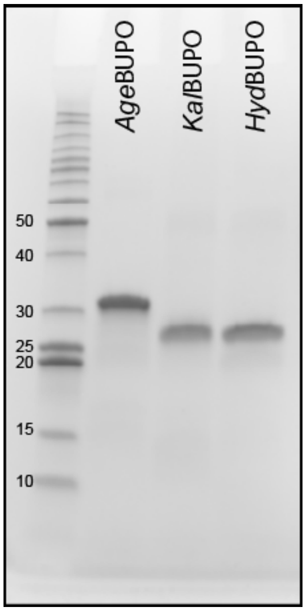


**Fig. S7. SDS-PAGE analysis of purified BUPOs.** The gel image shows the analysis of ~1 µg purified enzyme. The relevant sizes of the protein MW standard (Benchmark protein ladder, Thermo Fisher Scientific) are indicated in kDa next to the corresponding bands. The expected sizes for *Age*UPO, *Kal*BUPO and *Hyd*BUPO are 29.3 kDa, 23.8 kDa and 23.4 kDa, respectively, without considering the mass of heme. The gel was stained with Imperial Protein Stain (Thermo Fisher Scientific).

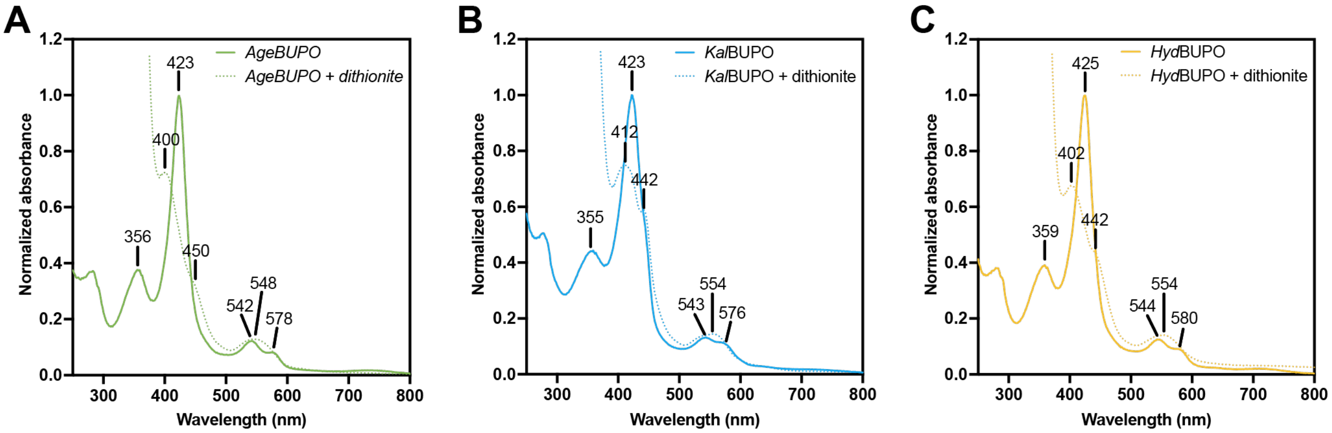


**Fig. S8. Spectrophotometric analysis of the resting state and reduced BUPOs.** The spectra of **(A)** *Age*BUPO, **(B)** *Kal*BUPO and **(C)** *Hyd*BUPO were recorded at 5-10 µM protein in 20 mM sodium phosphate buffer at pH 7.0, before (resting state; solid lines) and after the addition of sodium dithionite (reduced; dotted lines).

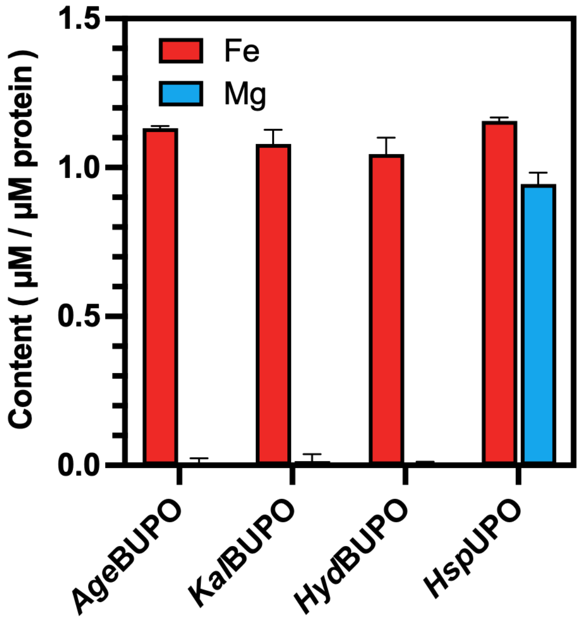


**Fig. S9. Iron and magnesium content in purified bacterial and fungal UPOs.** The values indicate the iron and magnesium concentrations in samples of pure protein measured by ICP-MS. The values were calculated by first subtracting the background iron/magnesium content measured in the buffer, followed by division by the concentration of heme-containing protein. The latter was estimated through the absorbance at 420 nm using the extinction coefficient of *Mro*UPO ($\varepsilon_{420}$ = 115 $mM^{-1}$ $cm^{-1}$) (23). The error bars indicate the standard deviation of the sample measurements (n=3), including the propagation of the standard deviation from the background measurements (n=3), $SD = (SD_1^2 + SD_2^2)^{1/2}$, and divided by the concentration of heme-containing protein. The *Hsp*UPO preparation used in this analysis was produced with the affinity tag StrepTagII to ensure high purity.

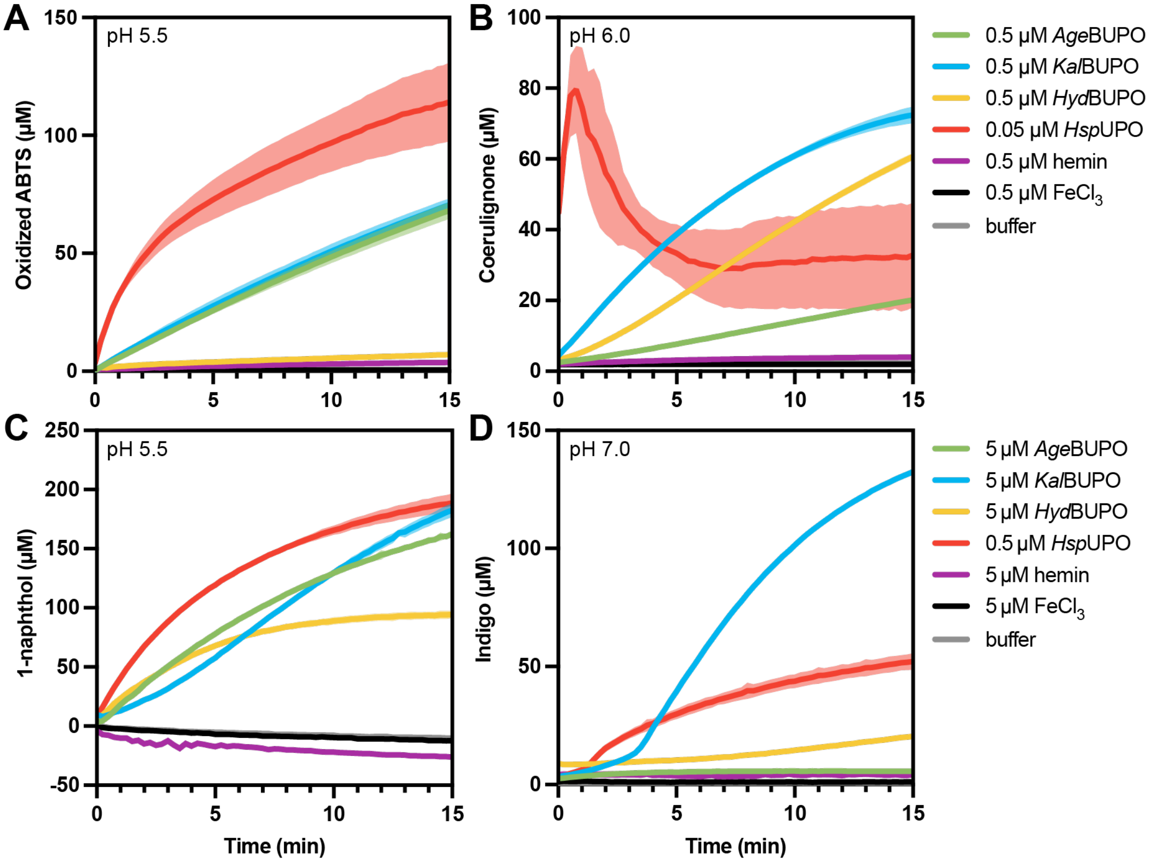


**Fig. S10. Time-course of peroxidase and peroxygenase activity by BUPOs and *Hsp*UPO.** Activity was investigated using four different spectrophotometric assays: **(A)** ABTS, 418 nm ($\varepsilon_{418}$ = 36 $mM^{-1}$ $cm^{-1}$ (24)); **(B)** 2,6-DMP, 469 nm ($\varepsilon_{469}$ = 27.5 $mM^{-1}$ $cm^{-1}$ (24)); **(C)** indole, 670 nm ($\varepsilon_{670}$ = 4.8 $mM^{-1}$ $cm^{-1}$ (24)); **(D)** naphthalene, 324 nm (quantified with a standard curve of 1-naphthol). The decrease in signal for the *Hsp*UPO trace in **(B)** is due to the precipitation of the reaction products (this reaction is very fast). Further details on the reaction conditions are provided in the **Supplementary M&M**. The peroxidase reactions on ABTS and 2,6-DMP were measured in 96-well plates using 0.5 µM enzyme, hemin chloride or $FeCl_3$ (0.05 µM for *Hsp*UPO), and the peroxygenase reactions on naphthalene and indole were measured using 5 µM enzyme, hemin chloride or $FeCl_3$ (0.5 µM for *Hsp*UPO). The formation of oxidized products was measured for 15 min. Note that *Hsp*UPO was used at 10-fold lower concentration than the rest of the enzymes in all reactions. The error envelopes indicate standard deviations between replicates (n=3).

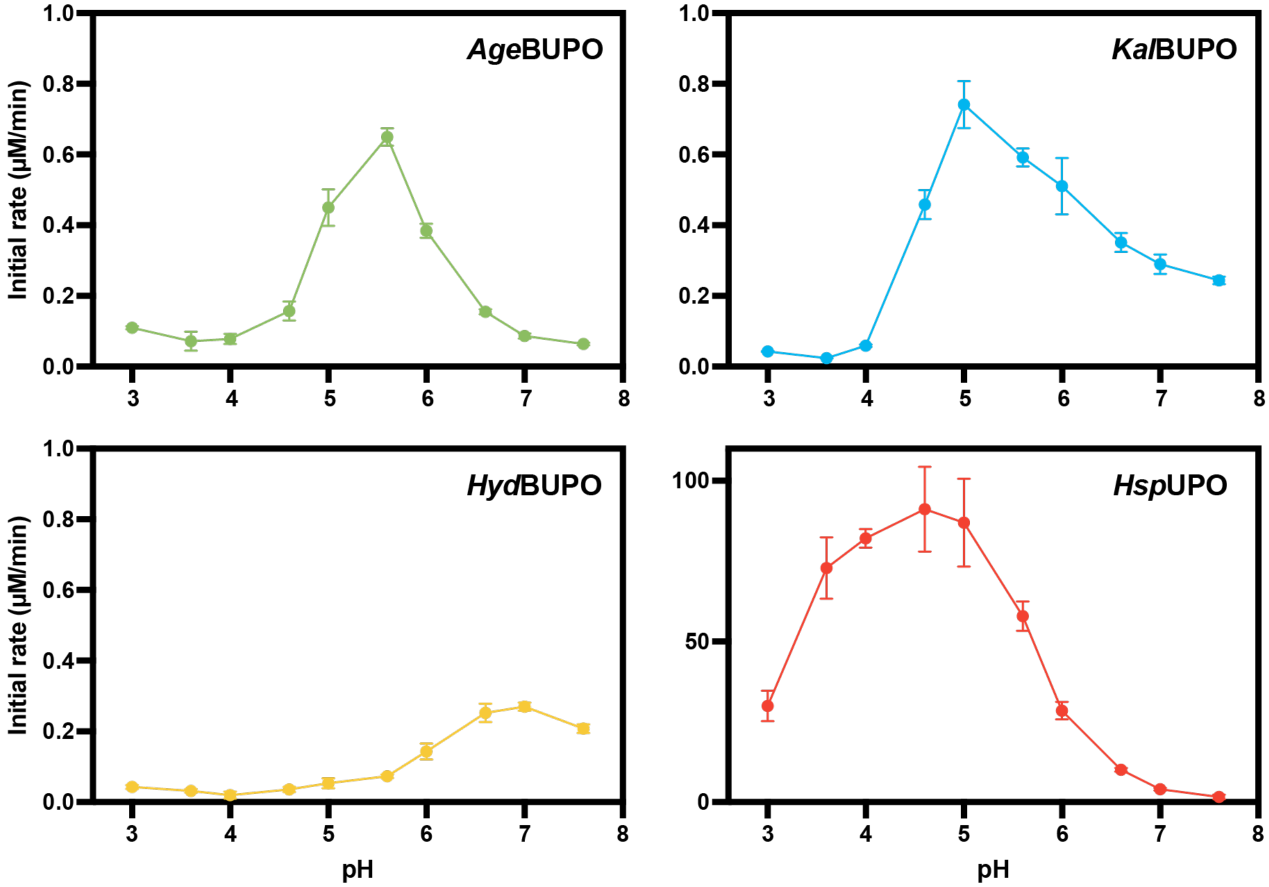


**Fig. S11. pH dependence of peroxidase activity by BUPOs and *Hsp*UPO**. Peroxidase activity was determined using an ABTS time-course assay, and initial rates were obtained from the linear portion of the progress curve. The reactions were carried out in McIlvaine (~75 mM) buffer at pH values ranging from 3.0 to 7.6, using 0.1 µM enzyme, 0.8 mM ABTS and 1 mM $H_2O_2$ in a total volume of 200 µL. The error bars indicate the standard deviation of replicates (n=3). Note that *Hsp*UPO is shown in a different scale than the rest of the enzymes.

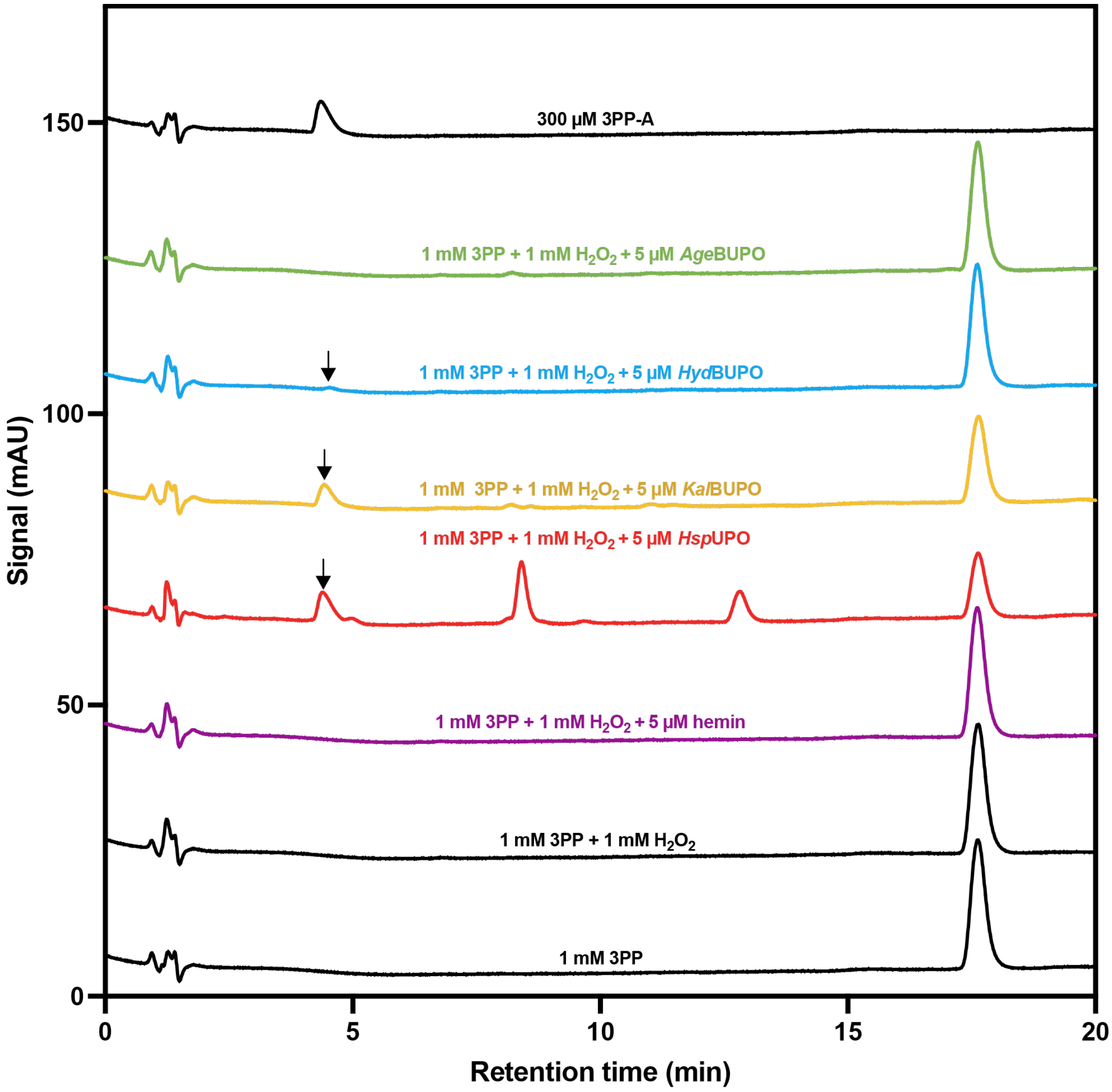


**Fig. S12. Enzymatic oxidation of 3-phenylpropanol (3-PP) by BUPOs and *Hsp*UPO monitored by HPLC.** The chromatograms show the reaction product 3-phenylpropanoic acid (3PP-A), indicated by arrows, eluting at approx. 4.5 min, whereas the substrate elutes at approx. 18 minutes. The peaks eluting between 4.5 and 18 min are other oxidized variants of 3PP (molecular identity not determined). Standards for 3PP and 3PP-A are shown at the lower and upper chromatograms, respectively. Control reactions with either buffer or hemin are shown in chromatograms two and three, counting from the bottom, respectively. The experiments were carried out in 50 mM sodium phosphate buffer, pH 6.0, at 30 °C for 1 hour.

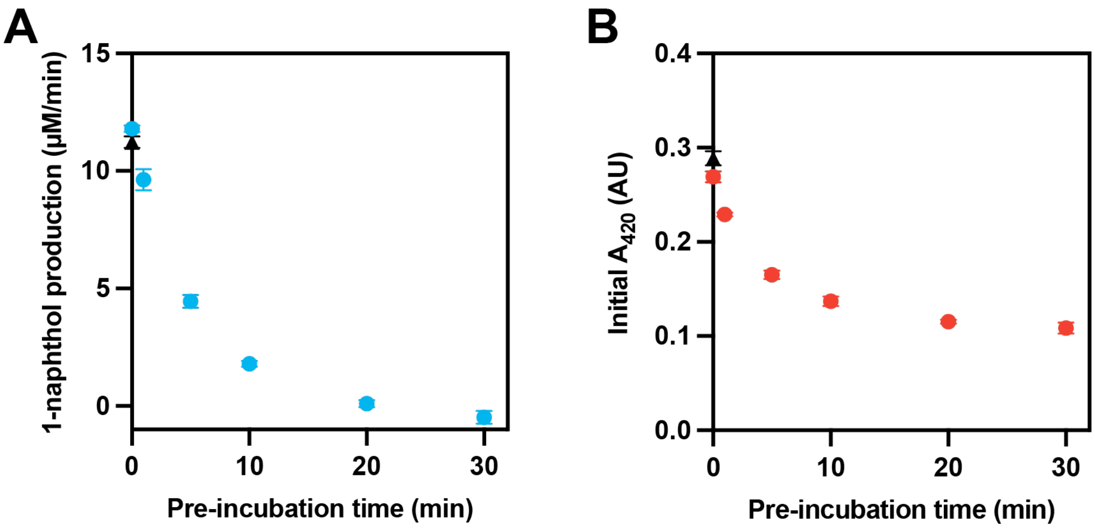


**Fig. S13. Inactivation of *Kal*BUPO by hydrogen peroxide.** The rate of 1-naphthol production **(A)** was measured at pH 5.5 in 100 mM sodium citrate buffer with 10 % acetone, 1 mM naphthalene, 1-1.25 mM $H_2O_2$ (1 mM freshly added + up to 0.25 mM potentially remaining from the pre-incubation stage) and 5 µM enzyme, in a final volume of 200 µL at room temperature. The enzyme (20 µM solution) had been subjected to pre-incubation in 50 µL of the same buffer containing 1 mM $H_2O_2$, at room temperature for 1-30 minutes, before the addition of 150 µL of naphthalene reaction mixture. The concentration of 1-naphthol was determined with a calibration curve of known concentrations and measuring the absorbance at 324 nm. The rates of product formation were derived from the initial linear phase of the reaction progress curve. Panel (**B**) shows the absorbance at 420 nm of the pre-incubated enzyme solutions, reflecting heme bleaching that occurred during the pre-incubation. Black triangles indicate the result for a reaction containing 1 mM $H_2O_2$, to account for possible effects of differences in $H_2O_2$ concentration due to $H_2O_2$ consumption at the pre-incubation stage. The error bars indicate the standard deviation of replicates (n=3).

**Table S1.** Fungal UPOs used in the phylogenetic analysis. The names of the two proteins used to first detect putative BUPOs appear **bold**.

| UniProt Accession | Name | Organism |
|---|---|---|
| A0A1L9X7H3 | *Aac*UPO | *Aspergillus aculeatus* |
| B9W4V6 | *Aae*UPO | *Agrocybe aegerita (Cyclocybe aegerita)* |
| A0A1L9U8A9 | *Abr*UPO-I | *Aspergillus brasiliensis* |
| A0A1L9UNI9 | *Abr*UPO-II | *Aspergillus brasiliensis* |
| A0A7R7WEQ5 | *Alu*UPO | *Aspergillus kawachii* |
| A2QLC7 | *Ani*UPO | *Aspergillus niger* |
| A0A7R8ALL8 | *Apu*UPO | *Aspergillus puulaauensis* |
| A0A100IPW8 | *Atu*UPO | *Aspergillus niger* |
| A0A4Q2DAL3 | *Cab*UPO-III | *Candolleomyces aberdarensis* |
| P04963 | *Cfu*CPO | *Caldariomyces fumago (Leptoxyphium fumago)* |
| A0A5C3KVD0 | *Cma*UPO-I | *Coprinopsis marcescibilis* |
| B9W4V8 | *Cra*UPO | *Coprinellus radians* |
| UPI002016A77D | *Cvi*UPO | *Collariella virescens (Achaetomiella virescens)* |
| A0A1Y2X6J4 | *Dsp*UPO | *Daldinia sp. EC12* |
| A0A067SWH8 | *Gma*UPO | *Galerina marginata* |
| A0A0C3CHS7 | *Hcy*UPO | *Hebeloma cylindrosporum h7* |
| A0A1Y2TH07 | ***Hsp*UPO** | *Hypoxylon sp. EC38* |
| A0A0C9Y789 | *Lam*UPO | *Laccaria amethystina LaAM-08-1* |
| A0A137QW42 | *Lsp*UPO-II | *Leucoagaricus sp. SymC.cos* |
| A0A9P7RSA8 | *Mor*UPO | *Marasmius oreades* |
| UPI002016A75C | ***Mro*UPO** | *Marasmius rotula* |
| A0A4Q4XVM9 | *Msp*UPO | *Monosporascus sp. 5C6A* |
| B2B6C6 | *Pan*UPO | *Podospora anserina (Pleurage anserina)* |
| A0A1W2TUZ6 | *Rne*UPO | *Rosellinia necatrix* |
| G2R8M6 | *Tte*UPO | *Thielavia terrestris* |

**Table S2.** Sequences of the proteins expressed in this study.

| Given name* | Code name (Fig. S6) | UniProt accession | Organism | Amino acid sequence† | SignalP prediction‡ | Supplier |
|---|---|---|---|---|---|---|
| | BU01-1 | A0A2E0HHX6 | *Pseudomonadales* bacterium | MSACGDASGPTREQLVAEFPTVERGSTRPENPEILCPFVRMLERSGLLDQTLAEQETLEVSTTELTAAADVFGCAPLECGTVAATVAVGQPGAAGVDIGRLHQAAGIAHDCGLTFAKGATQVTEARRQATLDRLALLADEQGRLTYPDLLEVKLATCAEEDVTITGAGRTETKLIFAYLGGVDNGYITLYDVESFLYASMPAVKTRYEVDLGLLSKVRAHHHHHH | OTHER | GenScript |
| | BU01-2 | A0A2E0HHX6 | *Pseudomonadales* bacterium | MKYLLPTAAAGLLLLAAQPAMAMKYLLPTAAAGLLLLAAQPAMAGDASGPTREQLVAEFPTVERGSTRPENPEILCPFVRMLERSGLLDQTLAEQETLEVSTTELTAAADVFGCAPLECGTVAATVAVGQPGAAGVDIGRLHQAAGIAHDCGLTFAKGATQVTEARRQATLDRLALLADEQGRLTYPDLLEVKLATCAEEDVTITGAGRTETKLIFAYLGGVDNGYITLYDVESFLYASMPAVKTRYEVDLGLLSKVRAHHHHHH | OTHER | GenScript |
| | BU01-3 | A0A2E0HHX6 | *Pseudomonadales* bacterium | MGDASGPTREQLVAEFPTVERGSTRPENPEILCPFVRMLERSGLLDQTLAEQETLEVSTTELTAAADVFGCAPLECGTVAATVAVGQPGAAGVDIGRLHQAAGIAHDCGLTFAKGATQVTEARRQATLDRLALLADEQGRLTYPDLLEVKLATCAEEDVTITGAGRTETKLIFAYLGGVDNGYITLYDVESFLYASMPAVKTRYEVDLGLLSKVRAHHHHHH | OTHER | GenScript |
| | BU02-1 | A0A4Q6C902 | Proteobacteria bacterium | MLIRSCLVALTGLSLSYYAFAAPGNPERLPSQIDPNISPAGFELSRAELVSRYSDIEEGSQQVENKKIVCPFLRMLERAGLFNPELETQSTLTVGIIKIASYAREFGCVVAGCGGVAAAVSAGQVTELASTPGKVNVEALHKALGISHECGLTFAKGGSVVDDATRDSTLAALKERADTLGRLTFDDLEAVKLSICEAQDVKISAPGRVEIGLIYTFLGGNERGFIDYDDVVRFFHAELPKTLGRPGIATHAHHHHHH | SP | GenScript |
| | BU02-2 | A0A4Q6C902 | Proteobacteria bacterium | MKYLLPTAAAGLLLLAAQPAMAMKYLLPTAAAGLLLLAAQPAMAAPGNPERLPSQIDPNISPAGFELSRAELVSRYSDIEEGSQQVENKKIVCPFLRMLERAGLFNPELETQSTLTVGIIKIASYAREFGCVVAGCGGVAAAVSAGQVTELASTPGKVNVEALHKALGISHECGLTFAKGGSVVDDATRDSTLAALKERADTLGRLTFDDLEAVKLSICEAQDVKISAPGRVEIGLIYTFLGGNERGFIDYDDVVRFFHAELPKTLGRPGIATHAHHHHHH | SP | GenScript |
| | BU02-3 | A0A4Q6C902 | Proteobacteria bacterium | MHHHHHHAPGNPERLPSQIDPNISPAGFELSRAELVSRYSDIEEGSQQVENKKIVCPFLRMLERAGLFNPELETQSTLTVGIIKIASYAREFGCVVAGCGGVAAAVSAGQVTELASTPGKVNVEALHKALGISHECGLTFAKGGSVVDDATRDSTLAALKERADTLGRLTFDDLEAVKLSICEAQDVKISAPGRVEIGLIYTFLGGNERGFIDYDDVVRFFHAELPKTLGRPGIATH | SP | GenScript |
| | BU03-1 | A0A160TBE6 | hydrothermal vent metagenome | MILLIRKLSLAAVFACSTLLVGCGGDVLSSEELVSLFPEVGPESTRAENTDILCPFQRMLKRSGLYDNAEDGEATSLKVKTGLASEAAEVFGCDKGSCGSIITLASIAQWNLGKLDLSRLHEAGSLLSHDCGLTFEFGGTTVSDSQRQFTLDRLLALANTEGQLQFDDLITVKQEICESQGVEMTVGGETEVKLIYAYLGGVERSFIDHSDVVRLLHATMPAYKTSAMVDLDLIGQVQAHHHHHH | LIPO | GenScript |
| | BU03-2 | A0A160TBE6 | hydrothermal vent metagenome | MKYLLPTAAAGLLLLAAQPAMAMKYLLPTAAAGLLLLAAQPAMAGGDVLSSEELVSLFPEVGPESTRAENTDILCPFQRMLKRSGLYDNAEDGEATSLKVKTGLASEAAEVFGCDKGSCGSIITLASIAQWNLGKLDLSRLHEAGSLLSHDCGLTFEFGGTTVSDSQRQFTLDRLLALANTEGQLQFDDLITVKQEICESQGVEMTVGGETEVKLIYAYLGGVERSFIDHSDVVRLLHATMPAYKTSAMVDLDLIGQVQAHHHHHH | LIPO | GenScript |
| | BU03-3 | A0A160TBE6 | hydrothermal vent metagenome | MHHHHHHGGDVLSSEELVSLFPEVGPESTRAENTDILCPFQRMLKRSGLYDNAEDGEATSLKVKTGLASEAAEVFGCDKGSCGSIITLASIAQWNLGKLDLSRLHEAGSLLSHDCGLTFEFGGTTVSDSQRQFTLDRLLALANTEGQLQFDDLITVKQEICESQGVEMTVGGETEVKLIYAYLGGVERSFIDHSDVVRLLHATMPAYKTSAMVDLDLIGQVQ | LIPO | GenScript |
| ***Age*BUPO (native)** | BU04-1 | A0A2W5V8F7 | *Archangium gephyra* | MKIQPSIPSVAPSSTTPAERVTSQPTPAAPKVEGFQAPRAAPATDLEGTSHGYVAPNKVESPFTNEEKVQLARKIPCPALAGAFNAGMLKVAKDGTVKIPDLERTLGQLGAGGLVTKVLTSAADATDDVKGSFNLFKLNGSNLDHTGSTGIRQNGVHPERFEKLMSFSKDGQRLTAKDLADAAESFAKEDPGLRGRITQQAELTAVLKIFGRTAEDGSKYFMRDDAKSLFVDGQIPASWEPPAVPGKKVGLGEVLGGTALGLFRQLVNGQAHHHHHH | OTHER | GenScript |
| *Age*BUPO (pelB-SP) | BU04-2 | A0A2W5V8F7 | *Archangium gephyra* | MKYLLPTAAAGLLLLAAQPAMAMKYLLPTAAAGLLLLAAQPAMASHGYVAPNKVESPFTNEEKVQLARKIPCPALAGAFNAGMLKVAKDGTVKIPDLERTLGQLGAGGLVTKVLTSAADATDDVKGSFNLFKLNGSNLDHTGSTGIRQNGVHPERFEKLMSFSKDGQRLTAKDLADAAESFAKEDPGLRGRITQQAELTAVLKIFGRTAEDGSKYFMRDDAKSLFVDGQIPASWEPPAVPGKKVGLGEVLGGTALGLFRQLVNGQAHHHHHH | OTHER | GenScript |
| *Age*BUPO (truncated) | BU04-3 | A0A2W5V8F7 | *Archangium gephyra* | MHHHHHHASHGYVAPNKVESPFTNEEKVQLARKIPCPALAGAFNAGMLKVAKDGTVKIPDLERTLGQLGAGGLVTKVLTSAADATDDVKGSFNLFKLNGSNLDHTGSTGIRQNGVHPERFEKLMSFSKDGQRLTAKDLADAAESFAKEDPGLRGRITQQAELTAVLKIFGRTAEDGSKYFMRDDAKSLFVDGQIPASWEPPAVPGKKVGLGEVLGGTALGLFRQLVNGQ | OTHER | GenScript |
| | BU05-1 | A0A0R3MHD1 | *Bradyrhizobium lablabi* | MSDHPVATAPGTALAGQFPVSPNNPCPFLRALVANGYVGGDVVPLSQISEIVGDASGQTGLGKMKVRIATWMVAVIANGLGPGRLFKSATSGAVLDQLRDGPLDKHGGGSRILDATAKVHEEQIDRLASFGKDCKDPAGGIETGLTAKEIETFMAANIKRDGDAARWYFPILMKGEWPVLLKILGKGEGEERYLSVAEVRTLFVERRLPKRIADRLPKPASPAHHHHHH | OTHER | GenScript |
| *Kal*BUPO (native) | BU06-1 | A0A2K9LIB1 | *Ketobacter alkanivorans* | MIIIKKSTWSAALLLVATNIFVAGCSDAPGPSREQLVAEYPSVEQGSTRQENIEIMCPFVRMLERSGLFDETFANQGDLDISTSELTSAAQEFGCAVLECGTVAATAAVGQPGGSGVDIERLHEAAGIAHDCGLTFEYGGTQVSDDRRDATITRLGELADEQGHLVYDDILQVKLETCDGEGVGITTAGRTETKLIFAYLGGVDNGFVTLDDVSSFLHAEMPSVKTRFMVDARQLGKVRAHHHHHH | LIPO | Twist Bioscience |
| ***Kal*BUPO (pelB-SP)** | BU06-2 | A0A2K9LIB1 | *Ketobacter alkanivorans* | MKYLLPTAAAGLLLLAAQPAMAMKYLLPTAAAGLLLLAAQPAMASDAPGPSREQLVAEYPSVEQGSTRQENIEIMCPFVRMLERSGLFDETFANQGDLDISTSELTSAAQEFGCAVLECGTVAATAAVGQPGGSGVDIERLHEAAGIAHDCGLTFEYGGTQVSDDRRDATITRLGELADEQGHLVYDDILQVKLETCDGEGVGITTAGRTETKLIFAYLGGVDNGFVTLDDVSSFLHAEMPSVKTRFMVDARQLGKVRAHHHHHH | LIPO | Twist Bioscience |
| *Kal*BUPO (truncated) | BU06-3 | A0A2K9LIB1 | *Ketobacter alkanivorans* | MHHHHHHASDAPGPSREQLVAEYPSVEQGSTRQENIEIMCPFVRMLERSGLFDETFANQGDLDISTSELTSAAQEFGCAVLECGTVAATAAVGQPGGSGVDIERLHEAAGIAHDCGLTFEYGGTQVSDDRRDATITRLGELADEQGHLVYDDILQVKLETCDGEGVGITTAGRTETKLIFAYLGGVDNGFVTLDDVSSFLHAEMPSVKTRFMVDARQLGKVR | LIPO | Twist Bioscience |
| | BU07-1 | A0A4Q6G3Q5 | Proteobacteria bacterium | MLERAGLYNKEVGMGGRLLVSISKITTLAKQWGCAIKECGLVATAVSAGQVTNLSTNPGFANLGALHRALGVSHECGFTFAKGGSVVSDEQRATSLSRLEALADSEGRLTFDNLNTVKKQICEEQGEVNTFASQVEVKLIYSFLGGKDRGFVDYDDVVRFFHAELPKTISAPSGLAHHHHHH | OTHER | Twist Bioscience |
| | BU08-1 | A0A4V2B2U3 | Pseudomonadota bacterium | MNPFKRLSTQLLCTLLLSSSAIAAPLNRAEFAEKFPQIEEGSSKPENKAIVCPFHRMLERAGLYDKEVGMGGRLLVSIAKITTLAKQWGCAIKECGTVATAVSAGQLTNLSTKPGFANLGALHRALGVSHECGFTFAKGGSVVSDEQRATSLSRLEALADSNGRLTFDNLNTVKNQICEEQGEVNTFASQVEVKLIYSFLGGKDRGFVDYDDVVRFFHAELPKTISAPSGL | SP | Twist Bioscience |

| | | | | | | |
|---|---|---|---|---|---|---|
| | BU08-2 | A0A4V2B2U3 | Pseudomonadota bacterium | MKYLLPTAAAGLLLLAAQPAMAMKYLLPTAAAGLLLLAAQPAMAAPLNRAEFAEKFPQIEEGSSKPENKAIVCPFHRMLERAGLYDKEVGMGGRLLVSIAKITTLAKQWGCAIKECGTVATAVSAGQLTNLSTKPGFANLGALHRALGVSHECGFTFAKGGSVVSDEQRATSLSRLEALADSNGRLTFDNLNTVKNQICEEQGEVNTFASQVEVKLIYSFLGGKDRGFVDYDDVVRFFHAELPKTISAPSGL | SP | Twist Bioscience |
| | BU08-3 | A0A4V2B2U3 | Pseudomonadota bacterium | MHHHHHHAPLNRAEFAEKFPQIEEGSSKPENKAIVCPFHRMLERAGLYDKEVGMGGRLLVSIAKITTLAKQWGCAIKECGTVATAVSAGQLTNLSTKPGFANLGALHRALGVSHECGFTFAKGGSVVSDEQRATSLSRLEALADSNGRLTFDNLNTVKNQICEEQGEVNTFASQVEVKLIYSFLGGKDRGFVDYDDVVRFFHAELPKTISAPSGL | SP | Twist Bioscience |
| | BU09-1 | M5DTC9 | *Thalassolituus oleivorans* MIL-1 | MILLIRKLSLAAVFACSTLLVGCGGDVLSSEELVSLFPEVGPESTRAENTDILCPFQRMLKRSGLYDNAEDGEATSLKVKTGLASEAAEVFGCDKGSCGSIITLASIAQWNLGKLDLSRLHEAGSLLSHDCGLTFEFGGTSVSDSQRQFTLDRLLALANTEGQLQFDDLITVKQEICESQGVEMTVGGETEVKLIYAYLGGVERSFIDHSDVVRLLHATMPAYKTSAMVDLDLIGQVQAHHHHHH | LIPO | Twist Bioscience |
| | BU09-2 | M5DTC9 | *Thalassolituus oleivorans* MIL-1 | MKYLLPTAAAGLLLLAAQPAMAMKYLLPTAAAGLLLLAAQPAMAGGDVLSSEELVSLFPEVGPESTRAENTDILCPFQRMLKRSGLYDNAEDGEATSLKVKTGLASEAAEVFGCDKGSCGSIITLASIAQWNLGKLDLSRLHEAGSLLSHDCGLTFEFGGTSVSDSQRQFTLDRLLALANTEGQLQFDDLITVKQEICESQGVEMTVGGETEVKLIYAYLGGVERSFIDHSDVVRLLHATMPAYKTSAMVDLDLIGQVQAHHHHHH | LIPO | Twist Bioscience |
| | BU09-3 | M5DTC9 | *Thalassolituus oleivorans* MIL-1 | MHHHHHHGGDVLSSEELVSLFPEVGPESTRAENTDILCPFQRMLKRSGLYDNAEDGEATSLKVKTGLASEAAEVFGCDKGSCGSIITLASIAQWNLGKLDLSRLHEAGSLLSHDCGLTFEFGGTSVSDSQRQFTLDRLLALANTEGQLQFDDLITVKQEICESQGVEMTVGGETEVKLIYAYLGGVERSFIDHSDVVRLLHATMPAYKTSAMVDLDLIGQVQ | LIPO | Twist Bioscience |
| | BU10-1 | A0A2D5VMJ4 | *Pseudomonas sp.* | MTFPRLSLSSLSSGITSHFARKAAAVLTLAVGMTGCGGDTYSADELAALYPQVAAGSTTPEDAEILCPFQRMIKRSGLLDDVLADGEFEVRNRLVTEASEIFGCASGACGTFVGYASLAQGNWNTLELNRLHEAGFLSHDCGLTFELGSITVDDSRRDFTLDRLTDLAVDGTLSLDNLMQVKQEICDLEGVEMTIGGETEVKLIYAYLGGSERGYVMNSDVSRFLHATLPAYKSSEYIDFSVVSEAHHHHHH | LIPO | Twist Bioscience |
| | BU10-2 | A0A2D5VMJ4 | *Pseudomonas sp.* | MKYLLPTAAAGLLLLAAQPAMAMKYLLPTAAAGLLLLAAQPAMAGGDTYSADELAALYPQVAAGSTTPEDAEILCPFQRMIKRSGLLDDVLADGEFEVRNRLVTEASEIFGCASGACGTFVGYASLAQGNWNTLELNRLHEAGFLSHDCGLTFELGSITVDDSRRDFTLDRLTDLAVDGTLSLDNLMQVKQEICDLEGVEMTIGGETEVKLIYAYLGGSERGYVMNSDVSRFLHATLPAYKSSEYIDFSVVSEAHHHHHH | LIPO | Twist Bioscience |
| | BU10-3 | A0A2D5VMJ4 | *Pseudomonas sp.* | MHHHHHHGGDTYSADELAALYPQVAAGSTTPEDAEILCPFQRMIKRSGLLDDVLADGEFEVRNRLVTEASEIFGCASGACGTFVGYASLAQGNWNTLELNRLHEAGFLSHDCGLTFELGSITVDDSRRDFTLDRLTDLAVDGTLSLDNLMQVKQEICDLEGVEMTIGGETEVKLIYAYLGGSERGYVMNSDVSRFLHATLPAYKSSEYIDFSVVSE | LIPO | Twist Bioscience |
| | BU11-1 | A0A7Z9PY58 | bacterium SCN 62-11 | MNIQTRLNTYTPAAHAMDSLPRTEQEQKPVPPSEDGKFTIDDIICPFQRVAYNEGVLKVDENGNATNLPEVLKEYAGAGWGLTKVANHAAKRLSTDGSHWQALWADSYNLQDLEGSSLDHKADTQILRGGFNQERLDKALSFSSDGERLTLEDLRRFQKSNLEEEPGRHGEIFGAAELALLVKVFGRTDGTGTKFISNKDFTTIFKDNKWPEGWEPPKAGSTNFLTVAQSVNEYFSMDDKIGEAKEAVASEAAPAKGSAKQACPFLSGQPFDLAEAAKQHSDRLEAHHHHHH | OTHER | Twist Bioscience |
| | BU11-2 | A0A7Z9PY58 | bacterium SCN 62-11 | MKYLLPTAAAGLLLLAAQPAMAMKYLLPTAAAGLLLLAAQPAMADSLDPAGLSDKLDNPKAALIPCPWWRTVINEDLVKVDGDGNVTMKDLRHALKATGVTFGLREGAILGVKRVAAQLAGQATGGITGFMHVLCMDKINVLDLPKSSLMHTGDSGTLRNGFNQENLERLLSFSSDGQRITANDLADANKKQVEADPGESGRKFGIAEYSILLNIFGRKDENGQKYLTKQDLTDVFKNNEFPENWEKPKVGFINLGKSIFGMFGRQKEETKAHHHHHH | OTHER | Twist Bioscience |
| | BU11-3 | A0A7Z9PY58 | bacterium SCN 62-11 | MHHHHHHADSLDPAGLSDKLDNPKAALIPCPWWRTVINEDLVKVDGDGNVTMKDLRHALKATGVTFGLREGAILGVKRVAAQLAGQATGGITGFMHVLCMDKINVLDLPKSSLMHTGDSGTLRNGFNQENLERLLSFSSDGQRITANDLADANKKQVEADPGESGRKFGIAEYSILLNIFGRKDENGQKYLTKQDLTDVFKNNEFPENWEKPKVGFINLGKSIFGMFGRQKEETK | OTHER | Twist Bioscience |
| ***Hyd*BUPO (native)** | BU12-1 | A0A1V3RV63 | *Hydrogenophaga sp.* A37 | MHHHHHHAPTTPRQKPVSPNNPCPFLRTLVAQGLVPDDVVPIGELTDAILKVARTGEGEPTLPAAAIRAVALAANGLGPLQLLRAGMDGVALNALRGGPLDKQGAGSGILSATATVDAEQLDRLDQFASDHVNARGRKERGLDRAALDRMMDANIERAASPRLLDRQLMDGEWPILLQVMGKEGKAGRYLSVKEVSDLFLHRRFPRRMAAALKDQRG | OTHER | Twist Bioscience |
| | BU13-1 | A0A2V8LIJ3 | Acidobacteria bacterium | MNTDPIQPTADERTSVVEKKATCPFIGSAVAQDALPIRNDANDPLAGIEDVRRLGNTGGGNLGDLLVFFASGNHAFMRGASGKLDAAVPLGFFSLDFPGSQGSHPGHSGILQGDPESLNSGRFSQADFDRLINLATDGFLKRSDVGRFIAENLIRDPKSKVLDHRTVALLAGDLVHIVESGFGFIGDLIKPNQADSHRDLEEKLTKLGGEDNLVGSSGEFGLLFAFFAHKPGSKTVAGEPALDIRDLKTMFVAKRLPEGWETWKKSRIDWVTNTTGLLISAANEYRKLKGTLRAGVAHHHHHH | OTHER | Twist Bioscience |
| ***Hsp*UPO§** | | A0A1Y2TH07 | *Hypoxylon sp.* EC38 | MRFPSIFTAVLFAASSALAAPVNTTTEDETAQIPAEAVIGYSDLEGDFDVAVLPFSASIAAKEEGVSLEKREAEAAPSPSSGWQAPGPNDVRAPCPMLNTLANHGFLPHDGKGITVNKTIDALGSALNIDANLSTLLFGFAATTNPQPNATFFDLDHLSRHNILEHDASLSRQDSYFGPADVFNEAVFNQTKSFWTGDIIDVQMAANARIVRLLTSNLTNPEYSLSDLGSAFSIGESAAYIGILGDKKSATVPKSWVEYLFENERLPYELGFKRPNDPFTTDDLGDLSTQIINAQHFPQSPGKVEKRGDTRCPYGYH | SP | Twist Bioscience |
| ***Hsp*UPO-StrepTagII§** | | A0A1Y2TH07 | *Hypoxylon sp.* EC38 | MRFPSIFTAVLFAASSALAAPVNTTTEDETAQIPAEAVIGYSDLEGDFDVAVLPFSASIAAKEEGVSLEKREAEAWSHPQFEKAPSPSSGWQAPGPNDVRAPCPMLNTLANHGFLPHDGKGITVNKTIDALGSALNIDANLSTLLFGFAATTNPQPNATFFDLDHLSRHNILEHDASLSRQDSYFGPADVFNEAVFNQTKSFWTGDIIDVQMAANARIVRLLTSNLTNPEYSLSDLGSAFSIGESAAYIGILGDKKSATVPKSWVEYLFENERLPYELGFKRPNDPFTTDDLGDLSTQIINAQHFPQSPGKVEKRGDTRCPYGYH | SP | |

* The names in **bold** correspond to the variants expressed in 1 L scale and purified (see main text).

† The portions of the sequences in orange are the native signal peptide (SignalP 6.0 (3)) or disordered region (in AlphaFold models) that were excluded in the truncated variants; predicted lipidation sites (SignalP 6.0 (3)) are highlighted in yellow (**C**) and were excluded in truncated variants; the added His-tag (or StrepTagII in case of *Hsp*UPO) is shown in red; the pelB (or alpha factor in case of *Hsp*UPO) signal peptide sequence is shown in blue.

‡ Result of signal peptide prediction using SignalP 6.0 (3), indicating the presence of a Sec/SPI signal peptide (**SP**), or a Sec/SPII lipoprotein signal peptide (**LIPO**). **OTHER** indicates the absence of signal peptide.

§ *Hsp*UPO is the fungal UPO used as reference in this study.

**Dataset S1 (separate file).** Complete list of the putative bacterial UPOs identified through FoldSeek and UniProt BLAST, including their UniProt and NCBI accession numbers, organism of origin, sequence length, isolation source and host (if available), SignalP prediction result (SP: Sec signal peptide (Sec/SPI); LIPO: Lipoprotein signal peptide (Sec/SPII); or OTHER: No signal peptide at all), and the clade in the phylogenetic tree they belong to.

**Dataset S2 (separate file).** List of proteins detected in the proteomic analysis of *Hydrogenophaga sp*. A37 and the quantification values for all biological replicates. *Hyd*BUPO (MGS5087929.1) is highlighted with red font and yellow background.
